# A CRITICAL ROLE FOR NEUTRAL SPHINGOMYELINASE-2 IN DOXORUBICIN-INDUCED CARDIOTOXICITY

**DOI:** 10.1101/2025.03.20.644150

**Authors:** Samia Mohammed, Victoria Franzi, Ya-Ping Jiang, Fabiola N. Velazquez, Monica E. Alexander, Folnetti A. Alvarez, Danielle Lambadis, Sam B. Chiappone, Anne G. Ostermeyer-Fay, Leiqing Zhang, Achraf A. Shamseddine, Daniel Canals, Ashley J. Snider, Richard Z. Lin, Yusuf A. Hannun, Christopher J. Clarke

## Abstract

Cardiotoxicity is a major side effect of Doxorubicin (Dox) that has hampered its clinical utility, and strategies to mitigate this cardiotoxicity are limited. Sphingolipids (SL) are central to the chemotherapy response in cancer but their role in normal tissue is less clear. Here, we identified the SL enzyme neutral sphingomyelinase-2 (nSMase2) as a critical mediator of chronic Dox-induced cardiotoxicity, establishing nSMase2 as a key downstream effector of Dox in cardiomyocytes (CM) and showing that *in vivo* loss of nSMase2 activity is protecting against chronic Dox-induced cardiac damage and dysfunction. Biologically, these studies link nSMase2 with Dox-induced CM senescence both *in vitro* and *in vivo* and identify the dual specificity phosphatase DUSP4 as a novel effector of nSMase2 in the Dox response. In addition to cementing a role for SL metabolism in Dox effects in normal tissue, this study advances nSMase2 as a target of interest for cardioprotection.

---

Although effective as a chemotherapeutic, the clinical utility of Doxorubicin is limited by the major side effect of cardiotoxicity (**1–3**). Considered to be largely irreversible, Dox-induced cardiotoxicity is typically associated with increased cardiomyocyte (CM) death, fibrosis, and disruption of myofibrillar architecture (**1–3, 4–6**). Current treatment options include β-blockers and ACE inhibitors but are typically used for symptom management and their effectiveness remains controversial (**7–9**). Dexrazoxane – the only FDA approved cardioprotective agent – also has limited clinical use owing to side effects (**10**), and concern that it can interfere with the anti-cancer activity of Dox (**11**). Consequently, a wider understanding of the pathogenesis of Dox-induced cardiac damage could establish novel targets for intervention.

Sphingolipids (SL) are a family of bioactive lipids implicated in various processes including growth arrest, senescence, and apoptosis (reviewed in **12, 13**). Ceramides (Cers) are the prototypical pro-death, anti-growth SLs (**14, 15**) and are produced through three major pathways: *de novo* synthesis, sphingomyelin (SM) hydrolysis by sphingomyelinases (SMases), and recycling of sphingosine derived from breakdown of complex SLs (salvage). While Cer accumulation is observed with multiple chemotherapeutics, the pathway of Cer production varies according to the specific chemotherapeutic and cell type (reviewed in **16**). However, irrespective of the source, neutralizing Cer levels by blocking production or increasing metabolism blunts chemotherapy-induced cell death and can underlie therapeutic resistance (**17–20**). Cer accumulation has also been implicated in various cardiac pathologies including ischemia reperfusion injury (**21**), acute myocardial infarction (**22**), heart failure (**23**), and cardiomyopathies (**24**). Dox treatment also increases Cer both in isolated CMs and *in vivo* in the heart (**25–28**) with studies suggesting that reducing Cer can alleviate Dox effects (**26–28**). While this raises the possibility of modulating cardiac SLs to mitigate Dox-induced cardiotoxicity, its effects on SL metabolism are broad – both in cancer and non-transformed cells – and Dox can regulate the SL network at multiple points (**29**). Thus, defining the key pathways of Dox-induced Cer production in the heart is critical for development of an appropriate intervention, particularly to avoid interfering with the anti-tumor activity of the drug

In this study, we identified nSMase2 as the major Dox-induced SL enzyme in CMs and find that nSMase2-activity null *fro/fro* mice were profoundly protected from chronic Dox-induced cardiac damage and loss of cardiac function. Biologically, nSMase2 was important for *in vitro* and *in vivo* Dox-induced CM senescence while discovery analysis identified the dual specificity phosphatase DUSP4 as a novel effector of nSMase2 and Cer that was important for Dox-induced senescence. Collectively, these results advance nSMase2 as a target of interest for cardioprotection.

## MATERIALS AND METHODS

### Materials

HL-1 murine CM were a gift from Dr. William C. Claycomb. AC16 human ventricular cardiomyoblasts and primary human CFs were from Millipore Sigma (St Louis, MO). Claycomb Medium, qualified fetal bovine serum (FBS) for HL-1 cells, L-Glutamine, Norepinephrine, Gelatin, Fibronectin, soybean trypsin inhibitor, anti-actin antibody, and doxorubicin-HCl (Dox) were from Millipore Sigma. RPMI, DMEM/F12 1:1 culture medium, fetal bovine serum (FBS), superscript III reverse transcriptase (RT), and penicillin/streptomycin solutions were from Life Technologies (Carlsbad, CA). Bio-Rad protein assay was from Bio-Rad (Hercules, CA). Antibodies for total PARP, p53, p21 were from Cell Signaling Technology (Danvers, MA). Antibodies for nSMase2 were from Santa Cruz Biotechnology (Santa Cruz, CA). HRP-conjugated secondary antibodies were from Jackson labs. Pierce chemiluminescence and Supersignal kits were from ThermoScientific (Suwanee, GA). Fluoroshield Mounting Media with DAPI was from Abcam (Cambridge, UK).

### Cell culture and siRNA

HL-1 cells were maintained on gelatin-fibronectin coated flasks in 10% FBS Claycomb media. AC16 cells were maintained in 12.5% FBS F12/DMEM containing 1% Pen Strep. Cell lines were cultured at 37°C, 5% CO_2_ in a humidified atmosphere and tested for mycoplasma contamination bi-monthly. MCF7 and 4T1 cells were maintained in 10% FBS RPMI; SKBR3 cells were maintained in 10% FBS F12/DMEM; MDA-MB-231 and E0771 cells were maintained in 10% FBS DMEM. For experiments with acute Dox treatment (24h of sustained treatment), cells were sub-cultured in 60mm (200-300K) or 100mm (600-800K) dishes and media was changed 1-2hr prior to treatment. For experiments with transient Dox treatment (1h Dox treatment followed by washout), cells were sub-cultured in 100mm dishes (200K). Cells were transfected with siRNA to a final concentration of 20nM using lipofectamine RNAimax (Life Technologies) according to manufacturer’s protocol. For acute Dox treatment, siRNA transfected cells were cultured for 48h prior to media change and Dox treatment. For transient Dox experiments, siRNA transfected cells were sub-cultured into 6-well trays (15-30K/well) and left for 24h before media change and Dox treatment. The siRNA used is in Supplemental Table 1 with AllStar siRNA (Qiagen) used as negative control.

### Animal care and model of chronic dox-induced cardiotoxicity

All experimental procedures used in this study were approved in advance by the Institutional Animal Care and Use Committee (IACUC) of Stony Brook University. Heterozygote breeders were obtained from the Sphingolipid Animal Cancer Pathobiology Core at Stony Brook University and were crossed to generate WT and *fro/fro* mice (**40** for characterization of *fro/fro* mice as nSMase2-activity null). For genotyping of mice, mouse tail DNA was used as template for PCR using a Fro WT-forward primer, a Fro mutant-forward primer, and a Fro reverse primer (all at 10μM). After 35 cycles of PCR, DNA was separated on a 2.5% agarose gel with SYBR-Safe. WT alleles run at 316 bp while the mutant allele runs at 358 bp (primer sequences in supplemental methods). For a chronic Dox-induced model of cardiotoxicity, dosing was based on previously published murine models of Dox-induced cardiotoxicity (**30–33**) and adjusted according to treatment tolerance of the specific mouse strain used in this study. Reported cumulative Dox doses in mouse cardiotoxicity models range from 15 to 40 mg/kg with repeated lower doses (such as 4 mg/kg) commonly employed to mimic clinical cardiomyopathy. To ensure manifestation of cardiotoxicity, initial studies targeted the highest reported cumulative dose (40 mg/kg) but this elicited high mortality. Accordingly, doses were adjusted to a cumulative 20-24 mg/kg dose to balance animal survival through the end of treatment while ensuring sufficient cardiac dysfunction. Thus, mice were administered either Dox (4mg/kg) or vehicle (sodium chloride 0.9% saline) via intraperitoneal injection (IP) twice per week, every other week up to a cumulative dose of 20-24mg/kg. Mice were subjected to echocardiography (see below) prior to treatment and 3-5 days after the final injection prior to euthanasia. Hearts were excised, weighed, sectioned, and either fixed in 10% formalin for paraffin embedding and immunohistochemistry (IHC), or snap frozen in liquid nitrogen for biochemical analysis. Frozen tissue was placed in long-term storage at -80°C prior to subsequent use.

### Echocardiography

For pre- and post-experiment echoes, mice were placed under isoflurane anesthesia and hair removed using commercially available cream. Cardiac function was measured by echocardiography using VEVO 3100 Ultra High Frequency ultrasound (FUJIFILM VisualSonics) with electrocardiograms (ECG) taken throughout. Parameters recorded include interventricular septal end diastole and end systole, left ventricular internal diameter end diastole and end systole, left ventricular posterior wall end diastole and end systole, Left Ventricle End Diastolic Volume (LVEDV), Left Ventricle End Systolic Volume (LVESV). The % Ejection Fraction (%EF) and % Fractional Shortening (%FS) were calculated from these measurements. Nine images per mice were averaged to obtain the final measurement.

### Analysis of cardiac troponin (cTNT) in plasma

At endpoint, plasma was isolated from chronic Dox-treated mice. For this, following isofluorane euthanasia, blood was drawn directly from the heart using needle and syringe into a microtube and placed on ice. To prevent clotting, acid citrate dextrose (ACD) solution was added to the syringe and microtube. The blood and ACD solution mixture was centrifuged (1300 RPM, 2m, 4C with no brake) and, once removed, the upper plasma layer (clear or pink) was collected and stored at -80°C. For subsequent use, plasma samples are thawed from -80°C on ice. 50μl of plasma was analyzed for cTNT levels using the Mouse cardiac troponin ELISA kit (CTNI-1-US; Life Diagnostics).

### Immunohistochemistry

Dissected hearts were placed in tissue cassettes and fixed in 10% formalin. After 24h, cassettes were submerged in 70% ethanol and submitted to the Stony Brook Research Histology Core for embedding, sectioning, and processing for hematoxylin & eosin (H&E) and Masson’s trichrome staining. For IHC analysis, unstained slides obtained from the core were baked in an oven overnight at 60°C, and then cooled at room temperature for 10 min before dewaxing and hydrating phases. For this, slides were immersed in xylene (3 x 5 min) and then placed in 5 different containers of ethanol ranging from 100% to 50% (5 mins each). Slides were subsequently immersed in distilled water (5 min) before placing in sodium periodate (0.005M, pH 2.5; 5 min). After a rinse (distilled water), slides were placed in sodium borohydrate (0.003M; 30 min). Slides were again rinsed (distilled water) before being placed in a pressure cooker with citrate buffer (10mM sodium citrate, 0.05% + 0.05% Tween-20, pH 6) and cooked (110-120°C, 10 min). After cooling, the slides were placed at 4°C for 30 minutes and rinsed with TTBS (Tween Tris-Buffered Solution; 1X TBS, 0.1% Tween). Slides were blocked (5% BSA-TTBS; room temp, 1h) and then incubated in primary antibody (4°C, overnight in a humid chamber). The next day, slides were washed (TTBS, 3 times, 5 mins on rocker), incubated in secondary (room temp, 1h), and washed (TTBS, 3 times, 5 mins on rocker). Following washes, DAB (Chromogen) was placed on each tissue sample for up to 5 minutes until color formation. To stop the DAB reaction, slides were placed in distilled water. Hematoxylin was added as counterstain (3 min), then slides were rinsed with fresh tap water and dehydrated again (4 ethanol baths ranging from 50%-100%, 2 mins each; 2 separate xylene baths for 2 min each). Finally, slides were mounted (Burkitt’s xylene based mounting media) and imaged the following day. For all IHC, a matched secondary only negative control was prepared in parallel.

### Protein extraction, protein precipitation, and immunoblot analysis

To extract cellular protein, cells were washed in cold PBS and directly scraped in RIPA buffer (50mM Tris, 150mM NaCl, 1% Triton X-100, 0.5%, 0.1% sodium dodecyl sulfate supplemented with protease inhibitor cocktail (#P-8340, Millipore Sigma) and phosphatase inhibitor cocktails 2 and 3 (#P5726 and #P0044, Millipore Sigma). Cells were further disrupted by sonication on ice (two 10s bursts at low intensity), and protein concentration was estimated by Bradford assay with Biorad reagent (#50000006, Biorad, Hercules, CA). Lysate aliquots were mixed with one-third volume of 4X Laemmli buffer (Invitrogen). For some analyses, protein precipitation with trichloroacetic acid (TCA) was performed. For this, equal volumes of cell lysate and 10% TCA solution were mixed and placed on ice for 30 min. Precipitated protein was pelleted by centrifugation (5 min, 1000rpm) and solubilized in 50μl 2% SDS, 10μl Tris, and 20μl 4X Laemmli buffer. All protein samples were vortexed for 2-3s, boiled for 5-10 min, and stored at -20C. Protein was separated on 4-20% SDS-PAGE gels using the Criterion system (Bio-Rad). Protein was transferred to nitrocellulose membrane by wet transfer in Tris-Glycine buffer (Bio-rad) containing 20% methanol. To block non-specific binding, membranes were incubated in 5% milk in 0.1% tween TBS for 1h at room temperature and rinsed in 0.1% tween TBS twice to remove excess milk. Primary antibodies were prepared in 2% bovine serum albumin in 0.1% tween TBS and membranes were incubated overnight at 4C. The next day, membranes were washed (3 x 10 min, 0.1% tween TBS, room temperature), incubated in secondary antibody for 45-60 min (1:5000 HRP-conjugated in 2% BSA in 0.1% tween TBS), and washed (3 x 10 min, 0.1% tween TBS, room temperature). To visualize bands, membranes were washed for 5 min in enhanced chemiluminescence (Pierce) prior to exposure to x-ray film. Information on the primary antibodies used can be found in Supplemental Table 2.

### Quantitative real-time RT-PCR (qRT-PCR) and SL gene PCR array

Analysis of gene expression by qRT-PCR using taqman assays was done as described previously (**29, 34**). Briefly, total mRNA was extracted using the PureLink RNA Mini Kit (Invitrogen) and converted to cDNA using the Superscript III supermix kit for real-time PCR (#11752-250; Life Technologies). PCR reactions were performed on the ABI 7500 real-time system using iTaq mastermix (Biorad) and Taqman assays (Life Technologies; see Supplemental Table 3 for list of Taqman assays used) with actin as reference gene. For the SL PCR array (custom from Life Technologies), reactions were run in 96-well plates with pre-coated taqman primers in each well using 10 μl of iTaq mastermix, 3 μl of cDNA, and 7 μl of water. Each array contained 2 sets of 48 genes, including 3 reference genes (18S, actin, GAPDH). For qRT-PCR samples, Ct values were converted to mean normalized expression using the ΔΔCt method with actin as reference gene. For the PCR array, results were converted to mean normalized expression using the ΔΔCt method with the arithmetic mean of actin, 18S, and GAPDH for normalization.

### Microarray analysis

For analysis of gene expression by microarray, total mRNA was extracted using the PureLink RNA Mini Kit (Invitrogen) and treated with DNase I to remove contaminating genomic DNA. Purified RNA was confirmed by bioanalyzer and analyzed using Clariom S arrays for human (Thermo Scientific) at the Stony Brook genomics core. For analysis, fold-change comparisons of target samples of control were performed and genes filtered at 2-fold up or down-regulation. For gene ontology analysis, nSMase2 specific upregulated hits were defined as those genes whose fold-change in Dox-treated AS siRNA samples was at least two-fold relative to vehicle-treated AS siRNA samples with an adjusted p-value of less than 0.05, but were not upregulated (fold-change less than two or adjusted p-value greater than 0.05) in Dox-treated nSMase2 siRNA samples, and unchanged in vehicle-treated nSMase2 siRNA samples. These hits were further analyzed by the web-based Enrichr program (**70**) for significant gene ontologies in the MSigDB Hallmarks database of gene ontologies. Fold-changes and p-values for the nSMase2 specific upregulated hits come from the R package *limma*. The p-values for the nSMase2 specific upregulated hits were adjusted according to the method of Benjamini-Hochberg, and the p-values of the gene ontologies were adjusted according to the default method of Enrichr.

### N-SMase enzymatic activity assay

*In vitro* analysis of neutral sphingomyelinase activity was performed by a mixed micelle assay as described previously using ^14^C-[methyl]sphingomyelin as a substrate (**34**).

### Lipid extraction and LC-MS/MS analysis

Following treatment, lipids were extracted by addition of 2ml 2:3 70% Isopropanol: Ethyl Acetate and stored at -80°C until submission to the Lipidomics Core at Stony Brook University for tandem liquid chromatography/mass spectrometry (LC/MS) as described previously (**29, 34**). For lipid analysis of mouse tissue, snap-frozen hearts were directly homogenized in 2ml of cell extraction buffer and submitted for analysis. For normalization of measured lipid values, total lipid phosphates of samples were analyzed as described previously (**29, 34**).

### Analysis of plasma membrane Cer

For this, the method previously described in Greene et al. (2023) was used (**35**). Briefly, following treatment of AC16 cells as indicated, cells were washed twice with serum free media (SFM), once with PBS, and fixed with 4% paraformaldehyde (20 min, room temp). Fixed cells were washed (SFM, x 2) and incubated with pseudomonas CDase (1h, 37 C). After treatment, cells were washed (SFM, x 2; PBS x 1) and immediately scraped in 2ml of cell extraction solvent (2:3 70% Isopropanol: ethyl acetate) containing internal standards. Extracted lipids (2ml) were transferred to glass tubes and an additional 2ml of cell extraction solvent was added. Samples were vortexed and centrifuged (3000 RPM, 10 min, room temp) with supernatants (approx. 4ml) being transferred to fresh glass tubes. An aliquot of extract (1ml) was removed to a separate glass tube for lipid phosphate analysis to normalize results. Both extracts were dried down under nitrogen and for lipidomic analysis, dried lipids (3ml) were resuspended in 150μl of mobile phase B (methanol containing 1mM ammonium formate, 0.2% formic acid) and transferred to appropriate vials. The remaining dried 1ml of extract was re-extracted and subject to lipid phosphate analysis as described previously (**29, 34, 35**). Results were normalized and expressed as pmol/nmol phosphate. From here, plasma membrane Cer was calculated as the sphingosine generated by the recombinant CDase as 1:1 molar ratio.

### Assessment of viable cell number by MTT and trypan blue exclusion

Following treatment, cells were washed once with warm PBS. Medium containing 5 mg/ml MTT (3-(4,5-Dimethylthiazol-2-yl)-2,5-Diphenyltetrazolium Bromide) was added for 30 min at normal cell culture conditions. The insoluble formazan product of MTT was dissolved with DMSO and quantified by measuring its absorbance at 570 nm using the SpectraMax plate reader. For trypan blue exclusion assays, cells were lightly trypsinized off dishes, pelleted, and resuspended in 1ml media. Equal volumes of cell suspension and trypan blue were mixed and incubated for 45-60s. Live and dead cells were counted under a light microscope using a hemocytometer to calculate the percent of dead cells.

### Analysis of Beta-galactosidase activity by staining

Senescence-associated beta-galactosidase activity was stained using a commercially available kit (#9860, Cell Signaling Technology). Following treatment, cells were washed twice with cold PBS, fixed for 10 mins (fixing solution), washed twice (cold PBS) and stained overnight at 37°C protected from light (staining solution). After 16-18h, staining solution was washed (cold PBS) and cells were imaged.

### Statistical analysis

Data were plotted using Graphpad. Comparison of two means was performed by unpaired student’s t-test, comparison of means of more than two groups were performed by 1-way ANOVA with appropriate post-test analysis, and comparison of experiments with two variables was by 2-way ANOVA with Bonferroni post-test analysis. For all statistical tests, a *p*-value of less than 0.05 was considered sufficient to reject the null hypothesis.

## RESULTS

### Dox increases in vivo and in vitro cardiac Cer levels and transcriptionally alters the sphingolipid network in CMs

To explore specific pathways of Cer production in the heart, we first assessed effects of chronic Dox treatment on heart SL levels (**Fig. 1A-C**). For this, mice received repeat injections of Dox (4mg/kg) or vehicle (DMSO in saline) up to a cumulative dose of 24mg/kg as before (**30–33**). Here, Dox robustly increased Cer (**Fig. 1A**) (2.43 ± 0.32 Dox vs 1.43 ± 0.04 Veh) with small, but significant, increases in Sph (0.010 ± 0.001 Dox vs. 0.007 ± 0.0002 Veh) and no changes in dhSph (0.028 ± 0.0005 Dox vs. 0.027 ± 0.0003 Veh) (**Fig. 1B).** There were also small but significant changes in S1P (0.042 ± 0.002 Dox vs. 0.033 ± 0.002 Veh) and dhS1P (0.020 ± 0.001 Dox vs. 0.015 ± 0.001 Veh) (**Fig. 1C**). The heart comprises multiple cell types with CMs and cardiac fibroblasts (CFs) as the two major components. To assess sites of Cer production, we treated human ventricular myocyte AC16 cells and primary human CFs with Dox (0.6 µM, 24h) prior to lipidomic analysis. Dox nearly doubled Cer levels in AC16 CMs but with minimal effect in CFs **(Fig. 1D)** although notably CFs had basally higher Cer levels than AC16 cells (data not shown). In contrast, Dox increased Sph in both AC16 and CFs (**Fig. 1E**) while S1P was suppressed in AC16 cells and unchanged in CFs. (**Fig. 1F**). Similar lipid changes were observed in mouse HL-1 CM cells (**Supp. Fig. 1A-1C**), and there was significant overlap in Dox-induced Cer species *in vitro* in AC16 cells and *in vivo* in the heart (**Supp Fig. 1D**). This suggests that CMs are the primary source of Dox-induced Cer accumulation in heart.

**Figure 1.**
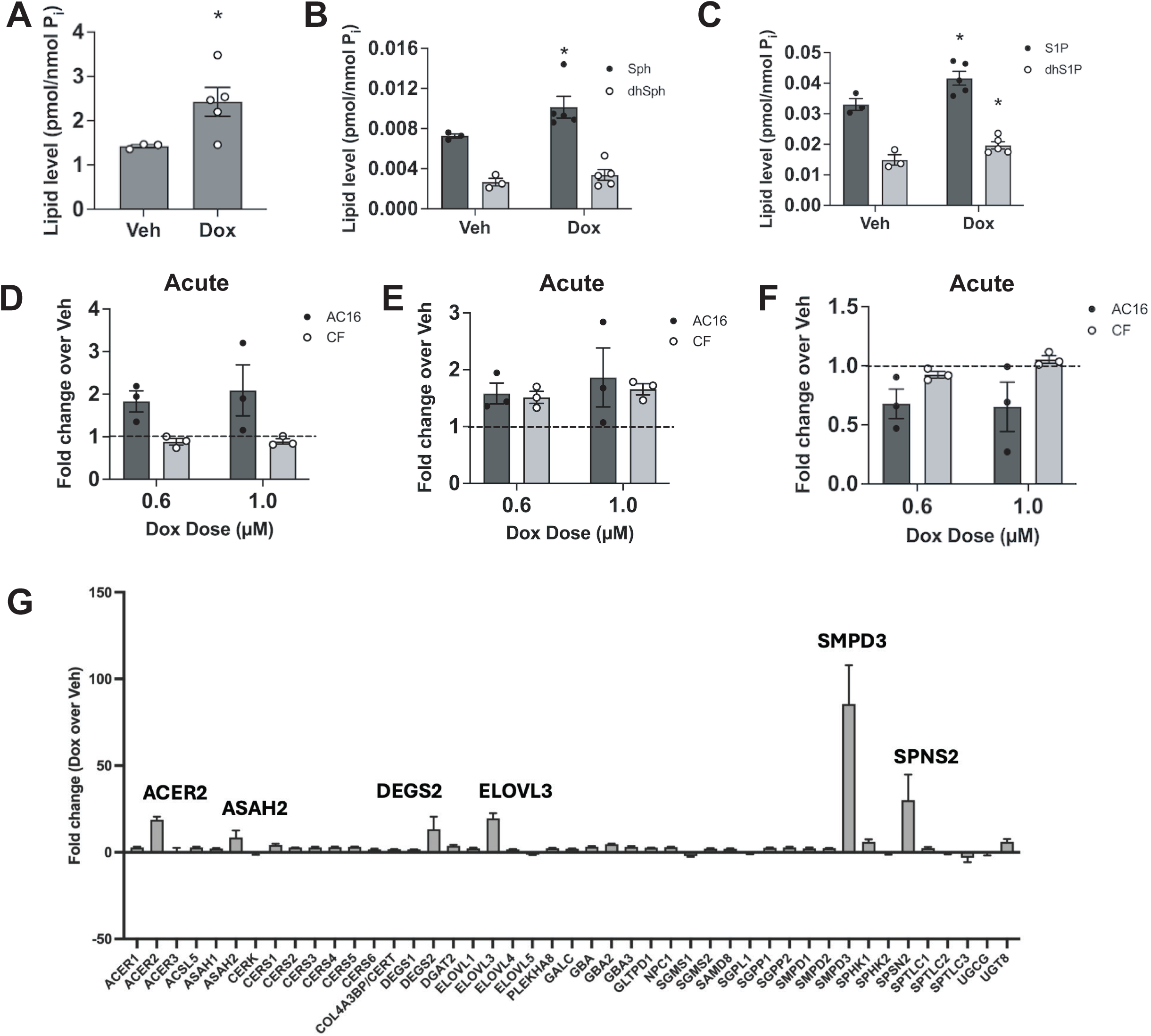
Doxorubicin induces Cer in vivo in the heart and in vitro in cardiomyocytes but not in cardiac fibroblasts. (A-C) Female FVB mice were treated with 6 doses of either vehicle or 4mg/kg Dox for up to 6 weeks. Lipids were extracted from heart tissues and analyzed by tandem LC/MS mass spectrometry for (A) Cer (* p<0.05, n = 3 Veh, 5 Dox; student’s t-test); (B) Sph and dhSph (* p<0.05, n = 3 Veh, 5 Dox; two way-ANOVA) (C) S1P and dhS1P (* p<0.05, n = 3 Veh, 5 Dox; two way-ANOVA). Data are expressed as mean ± SEM pmol/nmol lipid phosphate. (D-F) AC16 cardiomyocytes (AC16) or human cardiac fibroblasts (CF) were treated with vehicle or Dox doses as shown for 24h. Lipids were extracted and analyzed by tandem LC/MS mass spectrometry for (D) Cer; (E) Sph; (F) S1P. Data are expressed as mean ± SEM fold change lipid over Veh. Dashed lines represent vehicle control level (1.0-fold)(n = 3 biological replicates) (G) AC16 cells were treated with vehicle or 0.6μM Dox for 24h. RNA was extracted and analyzed by Taqman PCR array for the core sphingolipid genes shown. Data are expressed as mean ± SEM fold change Dox over Veh (n = 3 biological replicates)

### nSMase2 is induced by Dox in CMs and contributes to Cer generation

To probe which Cer generating pathways are involved in the Dox response, we performed PCR array analysis of the core genes in the SL network in Dox-treated AC16 cells (0.6μM, 24h) (**Fig. 1G**). While Dox increased expression of many SL genes (*ACER2, ASAH2, DEGS2, ELOVL3, SPNS2)*, the greatest induction was *SMPD3 (*85.5 ± 22.4-fold over vehicle). The *SMPD3* gene encodes for nSMase2, a key Cer-generating enzyme in response to cell stress (**34**). Extending these results, a Dox dose response (**Fig. 2A**) showed robust induction of nSMase2 at 0.4μM and 0.6μM with lesser induction at 1μM Dox. Similar effects were seen in mouse HL-1 CMs (**Supp. Fig. 2A**), human iPSC-derived CMs (**Supp. Fig. 2B**) and primary neonatal mouse CMs (**Fig. 2B**) although the extent of induction was more modest in these systems, and there were variances in the profile of nSMase2 induction. Importantly, only nSMase2 was significantly increased by Dox treatment (101-fold) (**Fig. 2A**) compared to nSMase1 (4.25-fold), nSMase3 (3.5-fold) and acid SMase (4-fold) (**Fig. 2C**) with similar results for nSMase1 and nSMase3 seen in HL-1 cells (**Supp. Fig 2C**) and iPSC-derived CMs (**Supp Fig. 2D**). Moreover, Dox treatment of CFs showed no effects on nSMase2 expression at lower doses although a modest (but significant) induction was seen at 1μM Dox (**Supp Fig. 2E**). Effects of Dox treatment on *in vitro* N-SMase activity showed robust increases at 0.4 µM (2.6-fold) and 0.6µM (2.5-fold) and a modest increase at 1.0 µM (1.48-fold) (**Fig. 2D**); it should be noted that the basal N-SMase activity reflects the sum of all N-SMases. Similar increases were measured in HL-1 cells treated with 0.6µM Dox (**Supp. Fig. 2F**). Furthermore, nSMase2 siRNA completely inhibited Dox-induced N-SMase activity in AC16 cells (**Fig. 2E, left**) consistent with effects of Dox and nSMase2 siRNA on nSMase2 protein levels (**Fig. 2E, right**).

**Figure 2.**
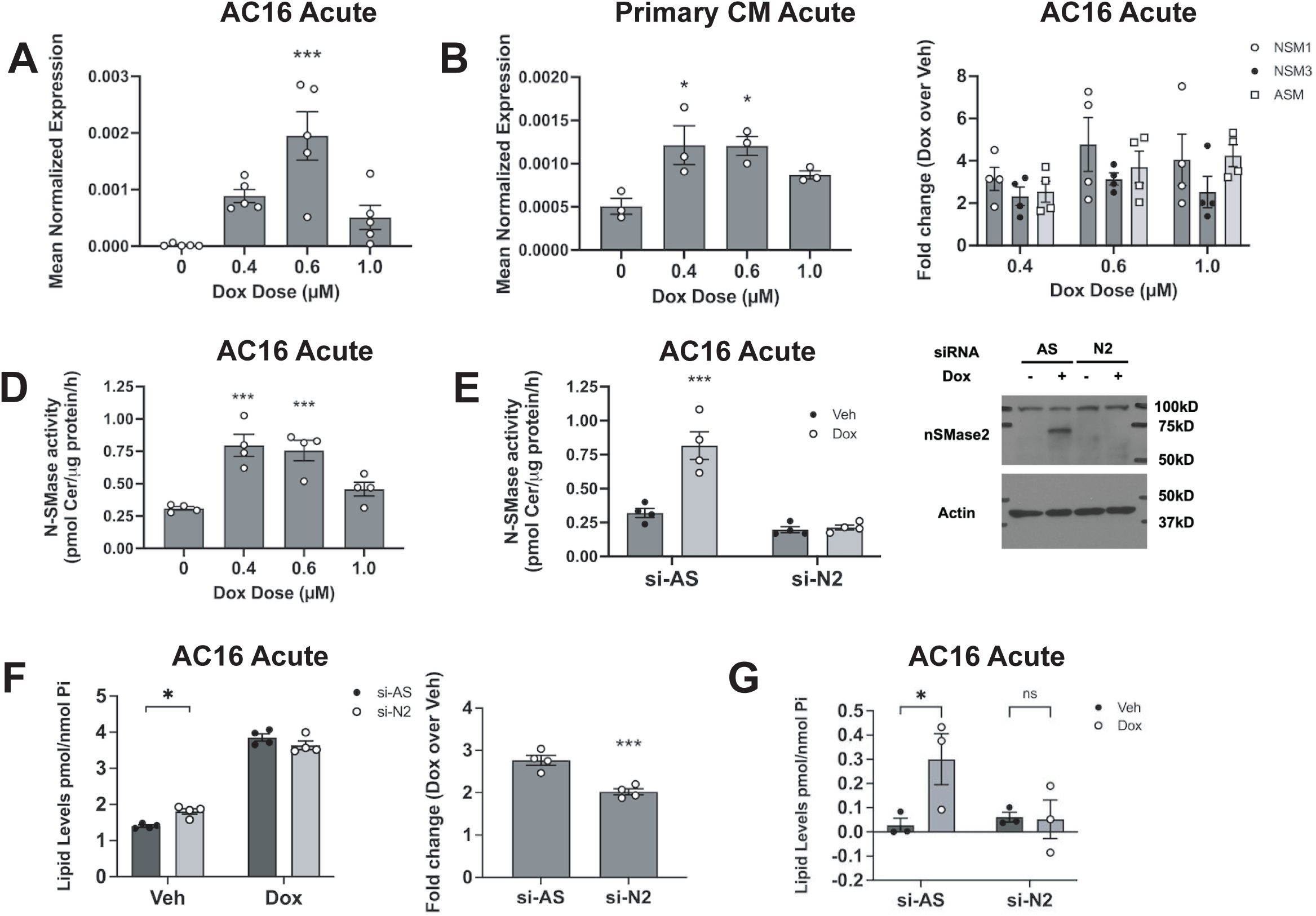
nSMase2 is induced by Dox in CMs and drives Cer accumulation at the plasma membrane. (A-C) Cells were treated with vehicle or the Dox doses as shown for 24h. RNA was extracted, converted to cDNA, and expression of nSMase2 (A, B) or the genes shown (C) was assessed by qRT-PCR using actin as reference gene. Data are expressed as mean ± SEM of mean normalized expression (A, B) ( *** p < 0.01, ** p < 0.02 vs vehicle; one way-ANOVA with Dunnet’s post-test. A: n = 5; B n = 3) or mean ± SEM fold-change over vehicle (C; n = 4). (D) Cells were treated with vehicle or the Dox doses as shown for 24h. *In vitro* N-SMase activity was assayed as described in ‘Materials and Methods”. Data are expressed as mean ± SEM N-SMase activity (*** p< 0.01 vs Veh; one way-ANOVA with Dunnet’s post-test, n = 4). (E-G) Cells were treated with negative control (si-AS) or nSMase2 (si-N2) siRNA for 48h prior to treatment with 0.6 μM Dox for 24h; (E) (left) In vitro N-SMase activity was assayed as described in ‘Materials and Methods”. Data are expressed as mean ± SEM N-SMase activity (*** p< 0.01 vs si-AS Veh; two way-ANOVA with post-test; n = 4); (right) Protein was extracted, precipitated, and analyzed by immunoblot for nSMase2 and actin as loading control. Representative of at least three biological replicates; (F)(left) Lipids were extracted and Cer levels measured by LC-MS. Data are expressed as mean ± SEM lipid level (* p <0.05, two-way ANOVA with post-test analysis, n = 4); (right) Data are expressed as mean ± SEM fold-change Dox over Veh (*** p < 0.01; student’s t-test, n = 4); (G) Plasma membrane Cer was analyzed as described in ‘Materials and Methods”. Data are expressed as mean ± SEM lipid level (* p <0.05, two-way ANOVA with post-test analysis; n = 3 biological replicates)

We next used siRNA to assess the contribution of nSMase2 to Dox-induced Cer levels. As before, Dox increased Cer levels in control cells yet while nSMase2 siRNA led to a small but significant increase in Cer levels in vehicle conditions (AS: 1.40 ± 0.03, N2: 1.80 ± 0.08, p<0.02), it had no significant effects with Dox treatment (**Fig. 2F, left**). Examining levels of individual Cer species revealed largely similar patterns with nSMase2 siRNA not impacting the Dox response (**Supp. Fig. 2G**). However, when considering fold-change of total Cer levels over Veh (i.e. taking increased basal levels into account), nSMase2 siRNA did reduce the Dox-specific response (AS: 2.76 ± 0.11, N2: 2.02 ± 0.07, p<0.01) (**Fig. 2F, right**). Nonetheless, this is consistent with prior studies implicating other pathways in Dox-induced Cer generation in CMs (**25–27**). Analysis of other lipids showed that nSMase2 knockdown led to a modest increase in Dox-induced Sph levels and a reduction in S1P levels in both Veh and Dox conditions (**Supp. Fig. 2H, I**). Finally, to specifically evaluate effects of nSMase2 induction on Cer, we utilized a recently developed assay for measurement of plasma membrane (PM) Cer (**35**). Here, Dox treatment strongly increased PM-Cer (**Fig. 2G**) and while nSMase2 siRNA slightly increased baseline PM-Cer, it completely prevented Dox effects. Taken together, this establishes a Dox-nSMase2-Cer pathway in CMs that is particularly prominent at the PM.

### Dox induction of nSMase2 is p53- and Top2B-dependent but independent of reactive oxygen species (ROS)

Dox has multi-faceted effects in CMs including ROS generation and activation of the DNA damage response (DDR) through Top2B and p53 (**36–39**). To assess where nSMase2 fits into these pathways, we first utilized the antioxidant N-acetylcysteine (0.5mM, 30 min pretreatment) to neutralize ROS prior to Dox (0.6μM, 24h) with results showing minimal effects (**Fig. 3A**). In contrast, siRNA knockdown of either Top2B (**Fig. 3B**) or p53 (**Fig. 3D**), but not Top2A (**Fig. 3C**) significantly reduced Dox-stimulated nSMase2 induction whereas combined Top2B and p53 siRNA treatment had modestly additive effects (**Fig. 3E**). Taken together, these results place nSMase2 as part of the Dox-induced DDR in CMs and link nSMase2 induction to key molecular mediators of Dox-induced cardiotoxicity.

**Figure 3.**
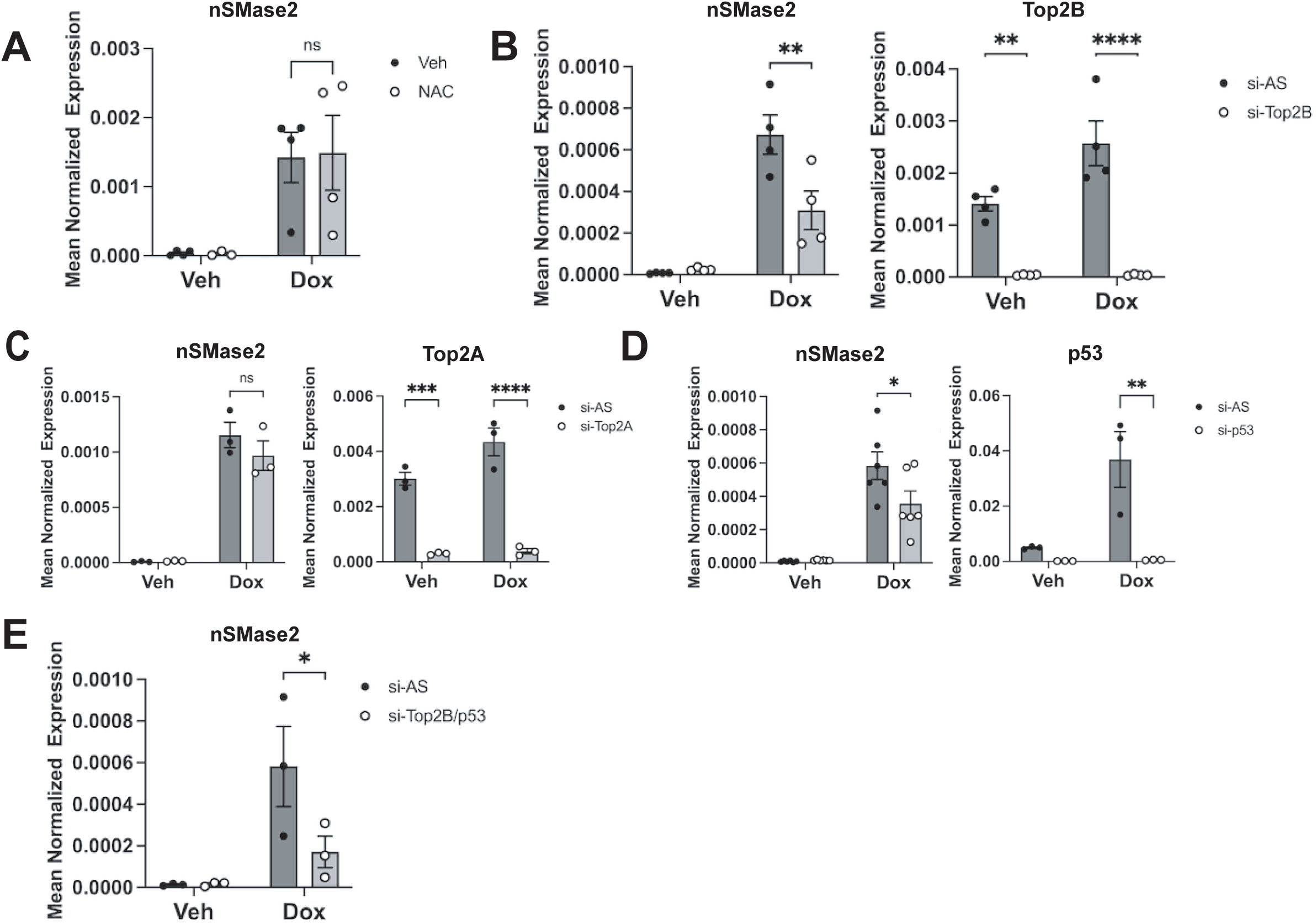
Dox induction of nSMase2 in AC16 CMs is dependent on Top2B and p53 but independent of reactive oxygen species and Top2A. (A) Cells were pre-treated for 30 mins with PBS or NAC (0.5mM) prior to treatment with vehicle (DMSO) or Dox (0.6μM, 24h)(Two-way ANOVA with post-test analysis; n = 4 biological replicates) (B) – (E) Cells were pre-treated for 48h with negative control (si-AS, 20nM) or target siRNA shown (20nM) prior to treatment with vehicle (DMSO) or Dox (0.6μM, 24h). (All panels) RNA was extracted, converted to cDNA, and gene expression for nSMase2, Top2B, Top2A, p53 as shown was assessed by qRT-PCR using actin as reference gene. Data are expressed as mean ± SEM of mean normalized expression; (B) **** p < 0.001, ** p < 0.02; two-way ANOVA with post—test analysis, n = 4; (C) **** p < 0.001, *** p < 0.01; two-way ANOVA with post-test analysis, n = 3); (D) ** p < 0.02, * p < 0.05; two-way ANOVA with post-test analysis, n = 3 (p53) or n = 5 (N2); (E) * p < 0.05, two-way ANOVA with post-test analysis, n = 3

### Loss of nSMase2 activity is protective against chronic Dox-induced cardiotoxicity in vivo

To evaluate the *in vivo* role of nSMase2 in Dox-induced cardiac dysfunction, we utilized *fro/fro* mice, which harbor a truncated catalytically inactive form of nSMase2 (**40**). This serves as an ideal loss-of-function model for nSMase2 activity while still allowing for expression of nSMase2 protein. Analysis of N-SMase activity in the brain showed substantial loss in the *fro/fro* strain compared to WT (WT: 100.0 ± 16.9%; fro/fro: 10.2 ± 1.0%) (**Fig. 4A**), consistent with prior results from nSMase2 knockout mice (**69–71**). Similarly, the N-SMase activity from hearts of *fro/fro* mice was significantly reduced compared to WT (WT: 100.0 ± 2.9%; *fro/fro*: 61.1 ± 3.1%) (**Fig. 4A**) although absolute N-SMase activity in heart was lower than in brain (data not shown). This supports a role for nSMase2 in the heart and is consistent with our *in vitro* activity data in CMs.

**Figure 4.**
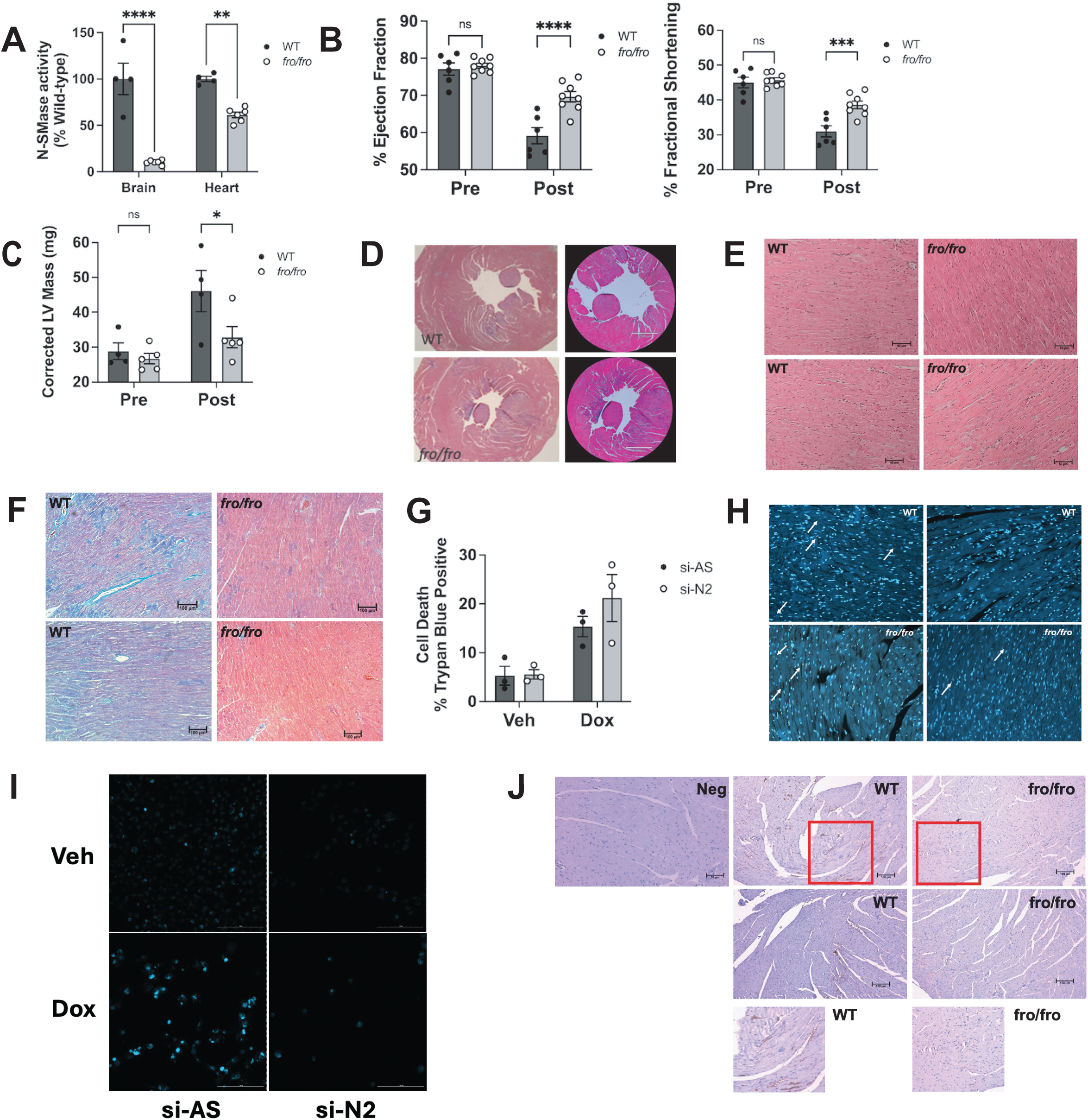
nSMase2 activity null mice (fro/fro) are protected from chronic Dox-induced cardiotoxicity. (A) *In vitro* neutral sphingomyelinase activity was assessed in brain and heart tissue from chronic Dox-treated female wild-type (WT) and fro/fro mice. Data are expressed as mean ± SEM % WT activity (** p<0.02; **** p<0.0001; two-way ANOVA with post-test analysis, n = 4 WT, n = 6 fro/fro) (B,C) Heart function was analyzed by echocardiography at baseline (pre) and 3-5 days post-chronic Dox treatment (Post) for female wild-type (WT) and fro/fro mice. Data are expressed as mean ± SEM for (B) Ejection fraction (EF); Fractional shortening (FS); and (C) Corrected left ventricular mass (*p<0.05; *** p<0.001; two-way ANOVA with post-test analysis, n = 6 WT, n = 8 fro/fro). (D) Transverse sections of hearts from female WT (top) and fro/fro mice (bottom) following chronic Dox treatment. Hearts were fixed, sectioned, and stained with H & E. Pictures were taken at 4 x magnification and are representative of at least three mice per group. (E) Heart sections from female WT (top) and fro/fro mice (bottom) following chronic Dox treatment stained with H & E. Pictures were taken at 10 x magnification and are representative of at least three mice per group. (F) Heart sections from female WT (top) and fro/fro mice (bottom) following chronic Dox treatment were subjected to Massons trichrome staining. Pictures were taken at 10 x magnification and are representative of at least three mice per group. (G) AC16 cells were pre-treated for 48h with negative control (si-AS, 20nM) or target siRNA (si-N2, 20nM) prior to treatment with vehicle (DMSO) or Dox (0.6μM, 24h). Cell viability was assessed by trypan blue exclusion assay. Data are expressed as mean ± SEM % trypan blue positive (n = 3 biological replicates). (H) Hearts from female WT (top) and fro/fro mice (bottom) following chronic Dox treatment were fixed, sectioned, and TUNEL stained. Images are from two separate mice per condition and are representative of at least four WT and four fro/fro mice. (I) AC16 cells were pre-treated for 48h with negative control (si-AS, 20nM) or target siRNA (si-N2, 20nM) prior to treatment with vehicle (DMSO) or Dox (0.6μM) for 1h. Treatment was washed out and cells cultured for 72h before staining for senescence-associated beta-galactosidase activity. Pictures representative of at least three biological replicates (J) Hearts from female WT (top) and fro/fro mice (bottom) following chronic Dox treatment were fixed, sectioned, and stained for p16/INK4a. Images are from two separate mice per condition and are representative of at least three WT and three fro/fro mice. Red box denotes inset area shown on right of figure.

To assess how the loss of nSMase2 activity affects cardiac function, echocardiography was used with emphasis on ejection fraction (%EF) and fractional shortening (%FS) as key cardiac parameters typically affected by Dox treatment (**1, 2**). No appreciable baseline cardiac phenotypes were observed in *fro/fro* mice compared to WT littermates. This was reflected in comparable baseline %EF (**Fig. 4B, left**) and %FS (**Fig. 4B, right**) results and suggests that nSMase2 is not required for normal heart function. After chronic Dox treatment (cumulative 20-24 mg/kg over five weeks), WT mice showed significant decreases in both %EF (59% vs 76% at baseline) and %FS (31% vs 45% at baseline). In contrast, *fro/fro* mice exhibited significant protections in cardiac function showing higher %EF (71%) and %FS (39%) following Dox treatment. Similarly, analysis of left ventricular mass (LVM) found no baseline differences (28mg WT vs 27mg *fro/fro*). However, while Dox treatment significantly increased LVM in WT mice (47mg), effects in *fro/fro* mice were more modest (34mg) (**Fig. 4C**). Cross-sections of heart tissue revealed an enlarged left-ventricle in Dox-treated WT mice consistent with a dilated cardiomyopathy while tissue sections from *fro/fro* mice showed a more typical phenotype (**Fig. 4D**). Histological analysis confirmed disorganization of myocardial fibers in WT mice (**Fig. 4E, left**) while their *fro/fro* counterparts showed more regular organization (**Fig. 4E, right**). Masson’s trichrome staining revealed fibrosis in WT hearts that was markedly reduced in fro/fro counterparts (**Fig. 4F**). Finally, analysis of plasma cardiac troponin (cTNT) – reflective of cardiac damage – revealed a marked reduction in cTNT in *fro/fro* mice compared to WT (**Supp. Fig. 3A**) although this was not statistically significant (p=0.11). Nonetheless, these results demonstrate that *fro/fro* mice are protected from chronic Dox-induced cardiotoxicity supporting nSMase2 as a crucial mediator of Dox effects in the heart.

### nSMase2 does not regulate Dox-induced CM cell death but is important for Dox-induced CM senescence

To understand how nSMase2 is involved in Dox-induced cardiotoxicity, we considered relevant biologies in Dox-treated CMs. Trypan blue exclusion analysis found that while acute Dox treatment (0.6μM, 24h) led to significant increases in CM cell death *in vitro*, there were no significant effects of nSMase2 knockdown (si-N2) compared to negative control siRNA (si-AS) (**Fig. 4G**) with similar results seen by MTT analysis of viable cell number (**Supp. Fig. 3B**). In line with *in vitro* results, TUNEL staining of heart sections from chronic Dox-treated mice (**Fig. 4H**) also showed little difference between WT and *fro/fro* hearts; however, it is also important to note that very few TUNEL positive cells were observed at all (typically 1-2 per field). As recent studies have begun to implicate CM senescence in Dox-induced cardiotoxicity (**41, 42**) and both Cer and SMase activity have previously been linked with senescence, (**43, 44**), we speculated that nSMase2 may have relevance in this context. To assess this, AC16 cells were treated with si-AS or si-N2 siRNA for 48h before transient treatment with vehicle or Dox (0.6μM, 1h; 3-5d incubation). In si-AS cells, Dox treatment induced a classic senescence morphology – enlarged, flat with a central nucleus – and increased senescence-associated beta galactosidase (Beta-gal) staining (**Fig. 4I, Supp Fig. 3C**). Remarkably, knockdown of nSMase2 mitigated Dox-induced senescence as shown by fewer cells with a senescent morphology and reduced Beta-gal staining (**Fig. 4I, Supp. Fig. 3C**). Importantly, similar results were observed with the pharmacological inhibitor GW4869 (**45**) which inhibited Dox-induced senescence (**Supp Fig. 3D**). To assess *in vivo* relevance, heart tissues were stained for p16/INK4a, which was previously reported as a marker for senescent CMs in Dox-treated hearts (**41**) (**Fig. 4J**). Here, hearts from Dox-treated WT mice showed clear patches of p16/INK4a staining suggestive of CM senescence while hearts from Dox-treated *fro/fro* mice showed little to no staining in hearts. Taken together, these data suggest that nSMase2 promotes Dox-induced cardiotoxicity through regulating CM senescence.

### DUSP4 is a downstream effector of nSMase2-Cer and regulates Dox-induced senescence

To explore nSMase2-dependent mechanisms, we performed a microarray in AC16 cells to identify Dox-induced genes that were affected by nSMase2 knockdown. Gene ontology analysis of those genes upregulated by Dox in control cells but significantly reduced in Dox-treated si-N2 cells (**Fig. 5A**) indicated involvement in TNF-alpha signaling, the p53 pathway, cholesterol homeostasis, IL-2/STAT5 signaling, and heme metabolism among others. An examination of these subsets identified the dual-specificity phosphatase DUSP4 as one of the strongest genes affected by nSMase2 knockdown (11.25 in si-AS vs 1.92-fold in si-N2) in the microarray with the related isoforms DUSP1 and DUSP6 also being induced and more modestly nSMase2-dependent (**Fig. 5B** for microarray expression data). Functionally, these genes were of interest as prior studies have implicated these DUSP isoforms in heart physiology and pathology (**46, 47**). Immunoblot analysis showed robust Dox induction of DUSP4 and DUSP1 in control si-AS cells although effects on DUSP6 were slight. Treatment with nSMase2 siRNA lead to a robust inhibition of DUSP4 induction but weaker effects on DUSP1 and DUSP6 as seen with the microarray (**Fig. 5C**); thus, we focused attention on DUSP4. Notably, the role of nSMase2 on DUSP4 was isoform specific as siRNA knockdown of nSMase1 did not significantly affect DUSP4 expression either basally or following Dox treatment (**Supp. Fig. 4C**). Finally, IHC analysis revealed robust DUSP4 staining in hearts of Dox-treated WT mice with a marked reduction in DUSP4 staining in Dox-treated *fro/fro* hearts (**Fig. 5D**). Taken together, this solidifies an nSMase2-DUSP4 connection in Dox-induced cardiotoxicity.

**Figure 5.**
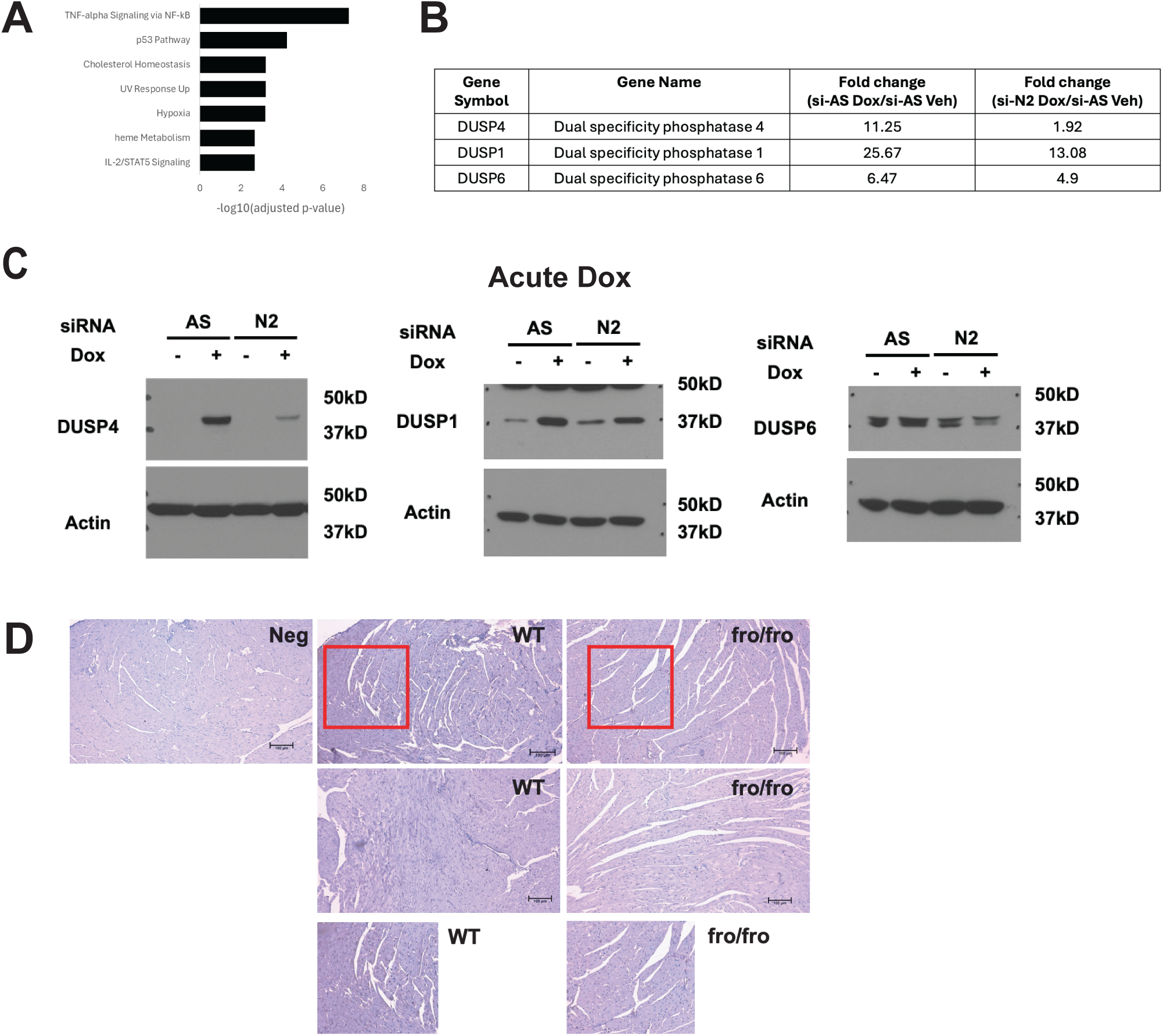
Dual specificity phosphatases are novel effectors of nSMase2. (A) Results of gene ontology analysis performed using Enrichr on microarray results. The negative log10 of the adjusted p-value for a test of enrichment performed by Enrichr is shown for the top gene ontologies enriched in the nSMase2 specific upregulated hits. (B) Microarray expression data for DUSP4, DUSP1, and DUSP6. Data are expressed as fold chance AS/Dox over AS/Veh (column 3) and N2/Dox over AS/Veh (column 4) – and are from analysis of 4 biological replicates. (C) AC16 cells were pre-treated for 48h with negative control (si-AS, 20nM) or target siRNA (si-N2, 20nM) prior to treatment with vehicle (DMSO) or Dox (0.6μM, 24h). Protein was extracted and analyzed for DUSP1, DUSP4, and DUSP6 by immunoblot using actin as loading control. Blots are representative of at least three biological replicates. (D) Hearts from female WT (top) and fro/fro mice (bottom) following chronic Dox treatment were fixed, sectioned, and stained for DUSP4. Images are from two separate mice per condition and are representative of at least three WT and fro/fro mice. Red box denotes the inset areas shown on right of figure.

These results prompted us to assess the relevance of the N2-DUSP4 connection to the effects of Dox (0.6μM, 1h before washout and 72h incubation). In control si-AS cells, Dox induced both nSMase2 and DUSP4 as seen with acute treatment while si-N2 blocked nSMase2 induction (**Fig. 6A**) and led to a significant decrease in DUSP4 mRNA and protein (**Fig. 6B**, **6C**). Importantly, pre-treatment with si-DUSP4 led to a reduction in SA-B-Gal staining following transient Dox treatment compared to controls (**Fig. 6D**). As the DUSP family are major regulators of MAPK signaling, we assessed effects of nSMase2 and DUSP4 knockdown on activation of ERK, JNK, and p38 MAPK following transient Dox treatment (0.6μM, 1h; washout and 72h incubation). In si-AS cells, Dox treatment led to increased phosphorylation of all three MAPK family members compared to vehicle. Notably, both nSMase2 and DUSP4 knockdown led to increased p-ERK signaling in Veh-treated cells but had no effect in the context of Dox treatment. DUSP4 siRNA also had no effect on p-JNK or p-p38 levels either in vehicle of Dox treated cells. In contrast, nSMase2 siRNA led to higher p-JNK and p-p38 levels in Veh-treated cells but did not increase levels further in Dox-treated cells. Thus, while DUSP4 appears to be connected with ERK signaling in AC16 cells, this may not be relevant in the context of Dox-induced (46, 47). Finally, treatment of AC16 cells with C6-Cer (5μM, 5 days) was sufficient to drive DUSP4 expression (**Fig. 6E**) and led to increased SA-B-Gal activity (**Fig. 6F**) as seen previously in fibroblasts (**43**, **44**) providing strong evidence of an nSMase2-Cer-DUSP4 connection that is important for Dox-induced senescence of CMs.

**Figure 6.**
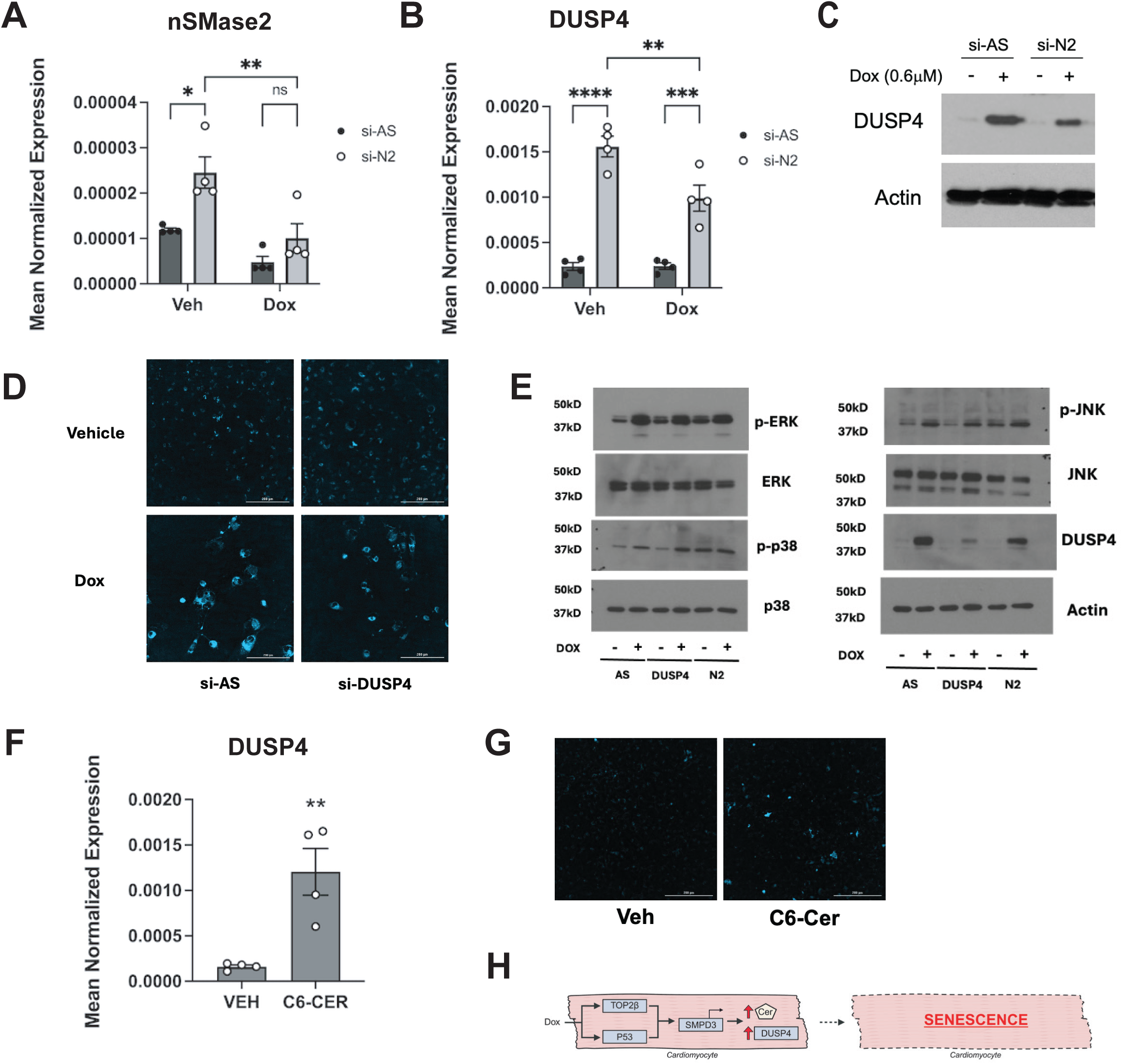
DUSP4 is a regulator of Dox-induced CM senescence. (A) AC16 cells were pre-treated for 48h with negative control (si-AS, 20nM) or target siRNA (si-N2, 20nM) prior to treatment with vehicle (DMSO) or Dox (0.6μM) for 1h. Treatment was washed out and cells cultured for 72h before RNA extraction, cDNA preparation and analysis of nSMase2 using actin as reference gene. Data are expressed as mean ± SEM of mean normalized expression (** P< 0.02, *** p< 0.01; two-way ANOVA with post-test analysis; n = 4 biological replicates). (B) Cells were treated as in A and analyzed for DUSP4 by qRT-PCR using actin as reference gene. Data are expressed as mean ± SEM of mean normalized expression (C) (** P< 0.02, *** p< 0.01, **** p<0.001; two-way ANOVA with post-test analysis; n = 4 biological replicates). (C) Cells were treated as in (A) and (B). Protein was extracted and analyzed by immunoblot for DUSP4 using actin as loading control. Immunoblot is representative of at least three biological replicates (D) AC16 cells were pre-treated for 48h with negative control (si-AS, 20nM) or target siRNA (si-DUSP4, 20nM) prior to treatment with vehicle (DMSO) or Dox (0.6μM) for 1h. Treatment was washed out and cells cultured for 72h before staining for senescence-associated beta-galactosidase activity. Pictures are representative of at least three biological replicates. (E) AC16 cells were pre-treated for 48h with negative control (si-AS, 20nM) or target siRNA (si-DUSP4 or si-N2, 20nM) prior to treatment with vehicle (DMSO) or Dox (0.6μM) for 1h. Treatment was washed out and cells cultured for 72h before protein was harvested and immunoblotted as shown. Pictures are representative of at least three biological replicates. (F) AC16 cells were treated with C6-Cer (5μM) for 5 days and analyzed for DUSP4 by qRT-PCR using actin as reference gene. Data are expressed as mean ± SEM of mean normalized expression (C) (** P< 0.02; student’s t-test, n = 4 biological replicates). (G) AC16 cells were treated as in (E). Cells were stained for senescence-associated beta-galactosidase activity. Pictures representative of at least three biological replicates. (H) Schematic of proposed pathway. Dox activation of Top2B and p53 signaling leads to upregulation of nSMase2, generation of Cer and increased DUSP4 expression. This promotes CM senescence.

### nSMase2 is dispensable for Dox-induced BC cell death

Given similarities in the Dox response of heart and cancer cells, we used a panel of human and murine BC cells to assess if nSMase2 was important for anti-cancer effects of Dox (**Supp. Fig 5A**). Here, Dox treatment led to significant decreases in viable cell number – as assessed by MTT assay – across all cell lines; importantly, pretreatment with GW4869 (10μM, 15 mins) had no effect. These results were broadly consistent with immunoblots for cleaved PARP as shown in MCF7, MDA, and SKBR3 cells (**Supp. Fig. 5B**) and suggests that, at least in BC, targeting nSMase2 does not interfere in anti-tumor effects of Dox.

## DISCUSSION

Multiple mechanisms underlying Dox-induced cardiotoxicity have been reported yet there continue to be limited treatments for mitigating this side effect without disrupting the anti-cancer activity of the drug. Accumulation of pro-death SLs such as Cer is central to the response to multiple chemotherapies across diverse cancers (**16**) and alterations in Cer production or metabolism have been associated with therapeutic resistance (**17–20**). While this increased interest in targeting the SL network to enhance therapeutic efficacy, utilizing this as a strategy to reduce chemotoxicities is less well explored.

Although Dox-induced Cer generation in the heart was reported some time ago, early studies into the relevant pathways and enzymes involved were hampered by a lack of molecular tools (**25–27**). Here, our primary finding implicates nSMase2 as a mediator of Dox-induced cardiotoxicity which was most strongly evidenced by *in vivo* studies showing that heart function was protected in the absence of nSMase2 activity, and these effects were associated with reduced cardiac damage, preserved myocardial architecture, and less fibrosis. This was complemented by *in vitro* studies identifying nSMase2 as the major Dox-induced SL enzyme in AC16 CMs and connecting nSMase2 induction with p53 and Top2β – both of which are established as components of the molecular machinery underlying Dox-induced cardiotoxicity (**37–39**). These results are consistent with prior reports utilizing the de novo SL synthesis inhibitor myriocin and showing that suppression of SL levels was protective against chronic Dox effects (**27**) and further supported by studies showing that protection from Dox-induced cardiotoxicity can be associated with reduced Cer levels (**28**). They also align with studies linking elevation of N-SMase activity and Cer with heart failure in various contexts (**21–24**), suggesting that nSMase2 may have a potentially broader role in heart pathologies. Intriguingly, targeting nSMase2 had only modest effects on total Cer levels *in vitro* but more substantially reduced PM-Cer (**Fig. 2**). While consistent with the multiplicity of Dox effects on the SL network noted previously (**29**), this suggests that PM-Cer may be a critical Cer pool for Dox-induced cardiotoxicity and that effects of Dox on Cer generated through other enzymes may not be as crucial. While this might seem to conflict with recent studies implicating CerS2 (**48, 49**), a possible scenario is that CerS2 is metabolically upstream – ultimately providing the SM substrate for nSMase2 and being directly linked with the same Cer pool; we are currently investigating this possibility. Nonetheless, beyond nSMase2, we identified other Dox-induced SL genes e.g. ACER2, SK1 (**Fig. 1G**) in AC16 cells further underscoring the potential for targeting SL metabolism in the heart, although confirming these effects are consistent across other CM relevant models e.g. iPSC-derived CMs, primary CMs (as done here for nSMase2) would be an important first step. An additional limitation of our study is that the protective effects observed here are in the acute post-treatment phase (c. 1 week post therapy) while the emergence of Dox-induced cardiotoxicity in clinical settings typically occurs 6-12 months post-treatment (**1–3**). Thus, to firmly establish nSMase2 as a target of interest for cardioprotection, future studies should assess if effects of nSMase2 loss persist at later time points (6-12 weeks post treatment). Additionally, our *in vivo* studies primarily utilized the *fro/fro* mouse and although this harbors an inactive nSMase2 protein as reported previously (**40**) and supported by results here (**Fig. 3A**), it nonetheless originated as a chemically induced mouse model of osteogenesis imperfecta. Thus, the possibility of other alterations that might impact the cardiotoxic Dox response cannot completely be ruled out. However, while preparing this manuscript, an independent study reported the nSMase2 inhibitor GW4869 was able to mitigate *in vivo* Dox-induced cardiotoxicity (**50**), providing additional supporting evidence for targeting nSMase2 for cardioprotection and consolidating our results with genetic approaches

Dox-induced cardiotoxicity is now appreciated to involve multiple cell types including CMs, CFs, vascular endothelial cells, and immune cells (**51–53**). However, studies here suggest that Dox effects on SL metabolism – or at least Cer accumulation – occurs primarily in CMs and not CFs. although it is important to emphasize that this is primarily based on *in vitro* cell studies and so may not fully reflect the *in vivo* situation. Nonetheless, significant overlap was observed between the Cer species increased *in vitro* in Dox-treated CM cell lines and *in vivo* in chronic Dox-treated hearts (**Supp. Fig. 1D**). Of note, our results showing no effects of Dox on Cer in CFs (**Fig. 1**) disagree with a prior study linking CerS2 with Dox-induced Cer generation in fibroblasts utilizing a comparable dose (0.7μM)(**48**). The reasons for this discrepancy are unclear but may plausibly be linked to cell type as the prior study utilized human foreskin fibroblasts as compared to the primary CFs that were utilized here. Furthermore, the effects on fibroblast Cer levels in this prior study were comparatively modest (30%) compared to what was observed in AC16 cells (close to 2-fold) (**Fig. 1D**). Biologically, nSMase2 was dispensable for *in vitro* Dox-induced CM cell death but was important for Dox-induced CM senescence, consistent with prior studies linking Cer and SMase with senescence in other cell types (**43, 44**). Notably, these results agree with recent studies implicating CM senescence as an important driver of Dox-induced cardiotoxicity (**41, 42**). While results with TUNEL would suggest that CM apoptosis may be less relevant with chronic Dox treatment, it is equally plausible that CM death occurs at proximal timepoints of Dox administration and may have concluded by the study endpoint used here. An additional point to note is that our studies primarily used immortalized AC16 cells as a CM model and although able to differentiate more towards the CM lineage in the presence of reduced serum (as used here), they may not be wholly reflective of adult CM biology. Nonetheless, the concordance between *in vitro* and *in vivo* systems provides strong supporting evidence of a role for nSMase2 in Dox-induced senescence. It should also be noted nSMase2 knockdown impacted a wide number of genes in Dox-treated AC16 cells (**Fig. 4A**) and, although subject to the same limitations of AC16 noted above, this does suggest that nSMase2 could have biological impacts in CMs beyond senescence. For example, nSMase2 inhibition was previously shown to prevent the effects of stretch-induced changes in the ventricular myocardium including blunted sympathetic responses (**54**). Such effects may be relevant as autonomic dysregulation can play a role in Dox-induced cardiac damage (**55, 56**) although changes indicative of altered autonomic function e.g. arrhythmias or major variability in average heart rate were not noted in either WT or fro/fro mice in our studies. Finally, it is also plausible that nSMase2 induction in CMs can drive changes in other cell types. Indeed, results showing reduced fibrosis in *fro/fro* hearts (**Fig. 4F**) suggest that *in vivo* effects of nSMase2 extend beyond the cell-intrinsic effects on CMs observed here. Possible mechanisms for this could be through secretion of inflammatory or pro-fibrotic mediators as, for example, nSMase2 may function upstream of S1P which can promote CF activation (**57**). NSMase2 is also a well-established regulator of EVs in multiple cell types (**58, 59**), and EVs have been linked with intercellular communication in the Dox-treated heart (**50, 60**). Still, as our studies utilized a global mouse model of nSMase2 activity loss, we cannot conclusively rule out roles for nSMase2 in other cell types relevant to the Dox response. To begin to address this, we have used αMHC-Cre mice to generate a conditional CM-specific nSMase2 knockout mice but studies with these mice proved to be problematic as we observed baseline defects both in knockout and Cre-expressing controls (data not shown). This is most likely due to adverse effects of Cre in the heart as has previously been reported (**61, 62**). Thus, we are in the process of generating inducible CM-specific nSMase2 knockouts to effectively answer this question.

Mechanistically, our studies identify DUSP4 as a primary and novel downstream effector of nSMase2 and as a regulator of Dox-induced CM senescence. This was further supported by C6-Cer effects on DUSP4 expression and senescence, as well as the reduced DUSP4 levels *in vivo* in chronic Dox-treated *fro/fro* mice. DUSP4 is a member of a family of proteins that are important negative regulators of MAPK signaling (reviewed in **63, 64**) and has previously been implicated in cardiomyopathy related to gene mutations in Lamin A/C (**65, 66**), suggesting a broader deleterious role in the heart. Consistent with this, depletion of DUSP4 in mice with the Lamin A/C mutation leads to improved cardiac function (**65, 66**). The connection of DUSP4 with senescence here is in line with prior studies linking DUSP4 with senescence of T-cells and breast cancer cells (**67, 68**). Thus, it would be of interest to determine if nSMase2-Cer induction may also be a factor in those contexts. Notably, the effects of DUSP4 in Dox-induced senescence here did not seem to be directly linked to regulation of MAPK signaling unlike the prior studies which involved suppression of ERK signaling (**67, 68**). One possible reason for this is that DUSP4 may regulate a specific subcellular pool of MAPK signaling akin to nSMase2 regulation of PM-Cer. It is also possible that the effects of DUSP4 during senescence are distinct from MAPK signaling and are mediated by effects on other pathways. For example, studies in breast cancer suggested that DUSP4 loss led to escape from the G1/S checkpoint through effects on cell cycle regulators (**68**). However, as these effects seemed to require concomitant p53 loss, their relevance for the Dox context is unclear. Finally, outside of the heart, Dox-induced DUSP4 expression was reported to play roles in drug resistance in breast cancer cells (**69**). Thus, targeting the nSMase2-DUSP4 axis may have benefits of both reducing cardiotoxicity and preventing the emergence of resistance.

In conclusion, this study provides novel insight into SL metabolism in the cardiac response to Dox, identifying nSMase2 as a critical mediator of Dox-induced cardiotoxicity and linking this to effects on CM senescence (**Fig. 6H**). We also establish DUSP4 as a novel downstream effector of nSMase2 and a new player in the cardiac Dox response. Efforts to mitigate Dox-induced cardiotoxicity have been hampered by broad concerns that they would interfere with the anti-tumor activity of Dox. Here, we show that targeting a key Cer-producing enzyme is sufficient to protect the heart without interfering with anti-cancer Dox activity as other pathways of Cer production remain intact. Overall, this provides a strong base of evidence to advance nSMase2 as a target of interest for cardioprotection.

## ACKNOWLEDGEMENTS

The authors would like to acknowledge the technical support provided by the Research Histology Core Laboratory - Department of Pathology, the Cancer Center Lipidomics Shared Resource, the Stony Brook Genomics Core, the Sphingolipid Animal Pathology Core, and the Division of Laboratory Animal Resources at Stony Brook University. We thank the Damaghi lab at Stony Brook University for use of and assistance with the Cytation imaging microscope. We would like to thank Andrew Resnick for assistance with figures and are also grateful to all members of the Lipid Cancer lab at Stony Brook University for helpful feedback and discussion

## DATA AVAILABILITY

The data that support the findings of this study are available from the corresponding author upon reasonable request.

