## Supplemental Information for "A CRITICAL ROLE FOR NEUTRAL SPHINGOMYELINASE-2 IN DOXORUBICIN-INDUCED CARDIOTOXICITY"

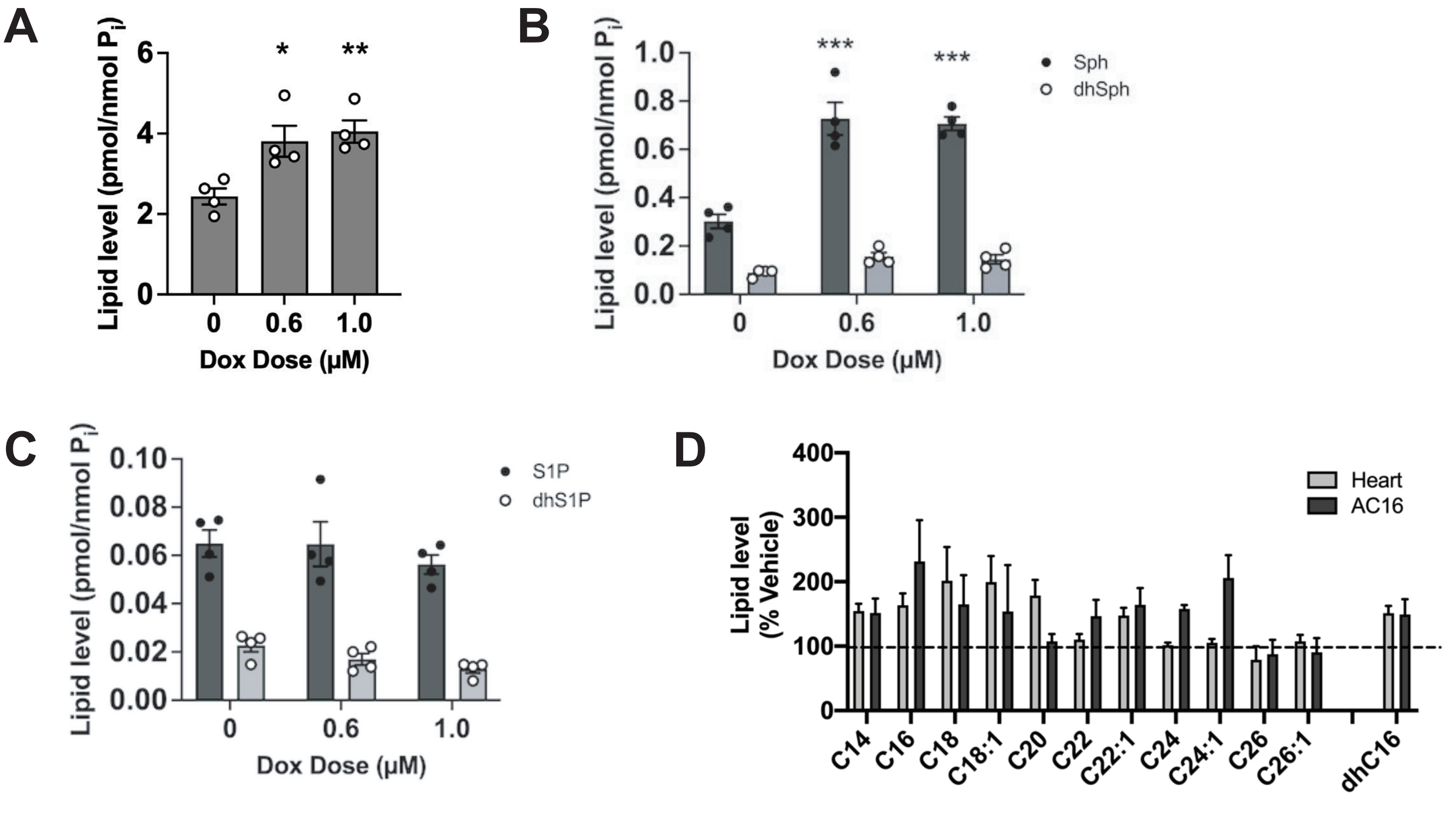


### SUPPLEMENTAL FIGURE 1

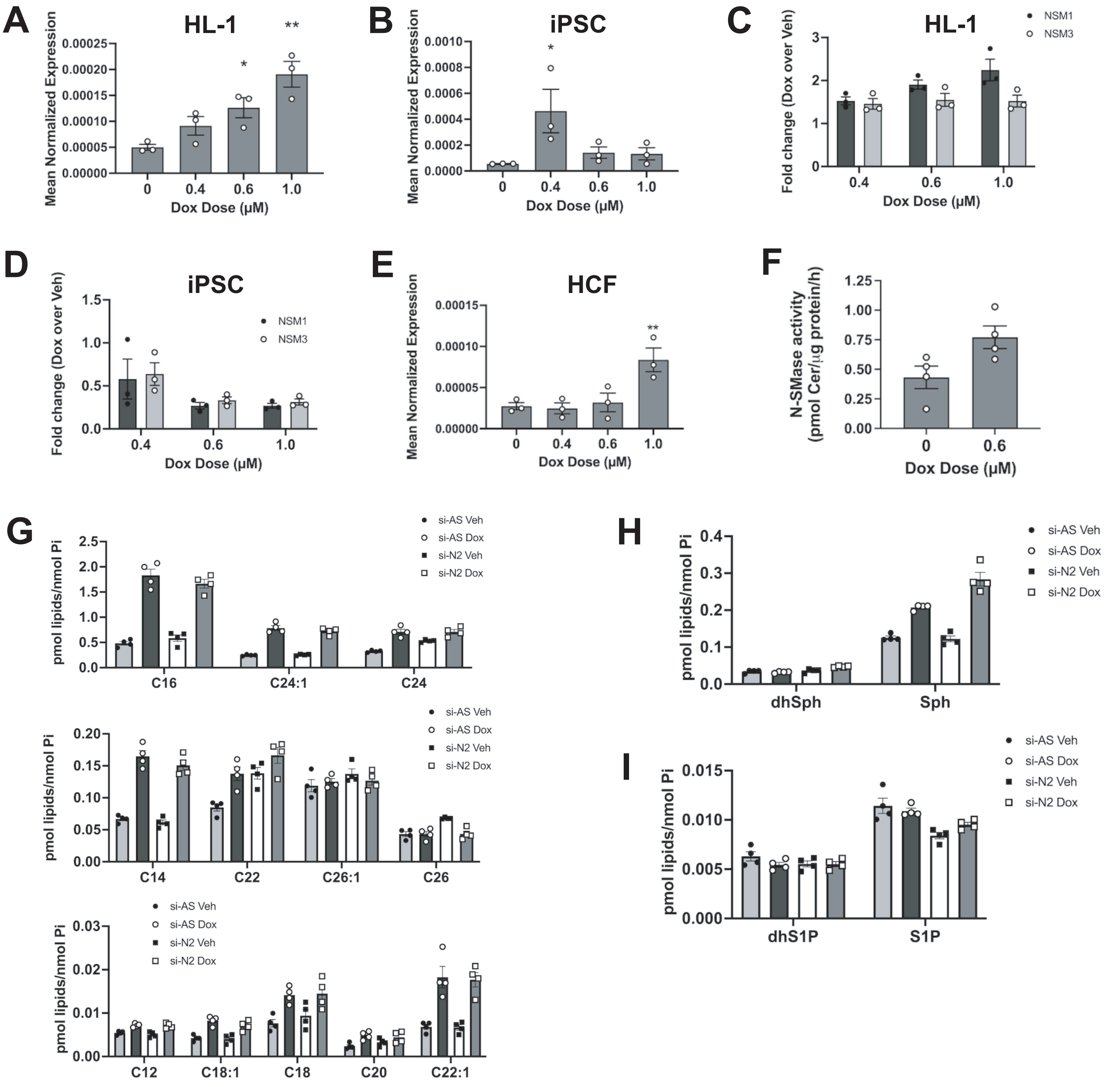


### SUPPLEMENTAL FIGURE 2

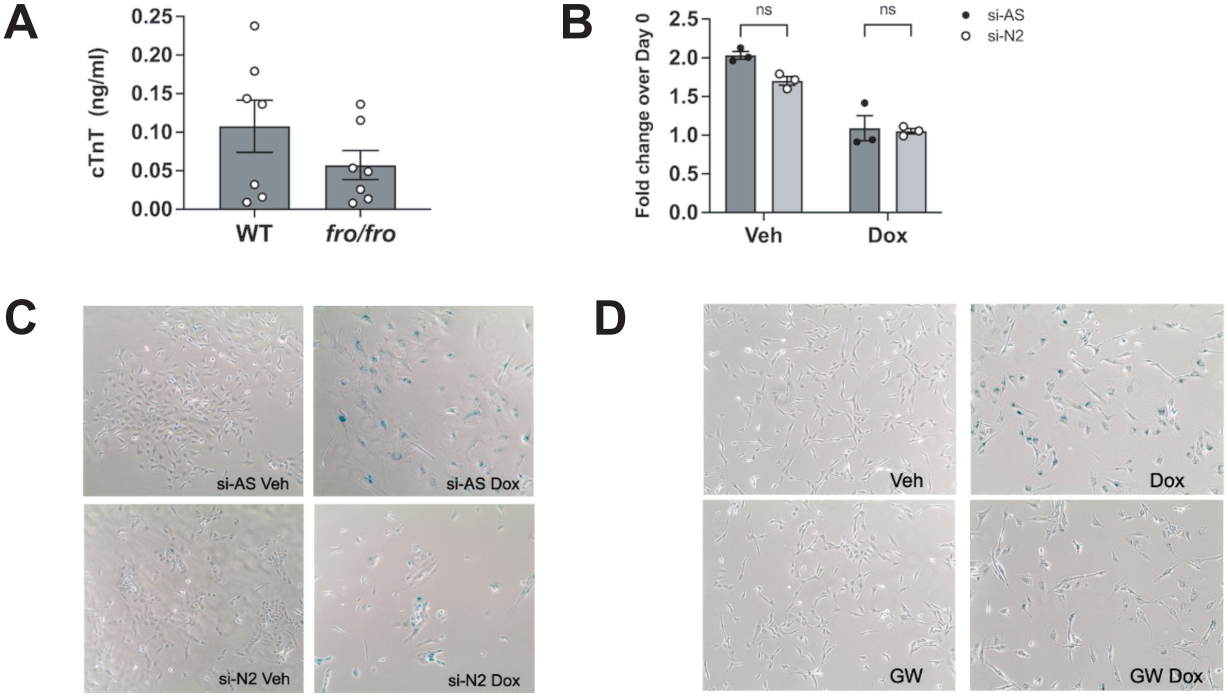


### SUPPLEMENTAL FIGURE 3

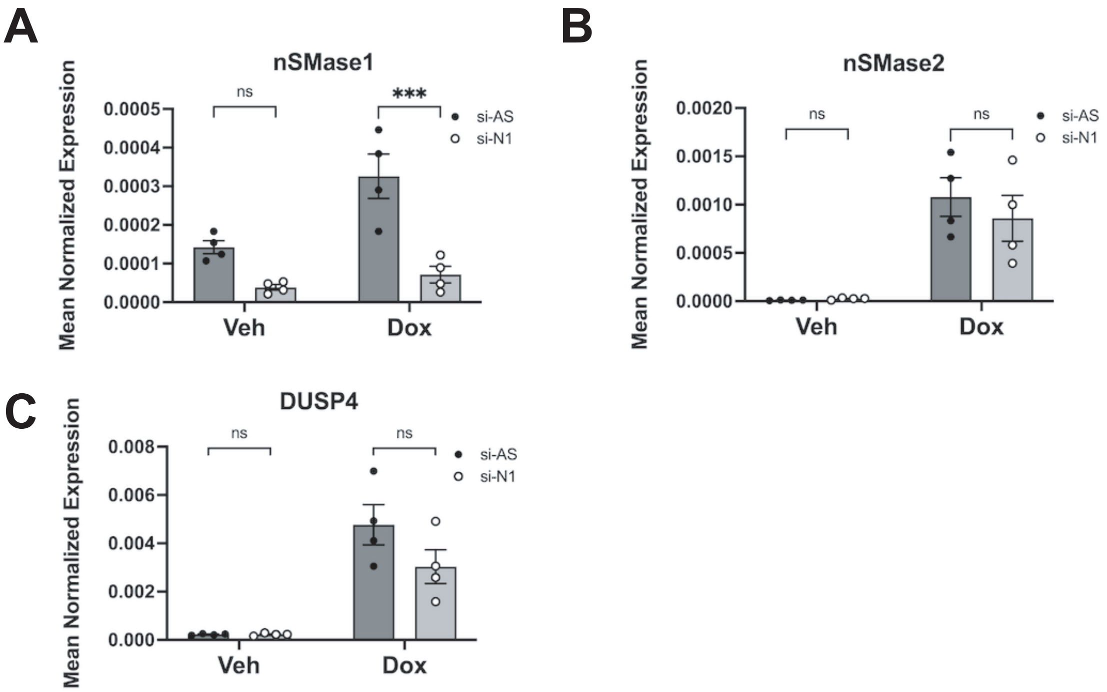


### SUPPLEMENTAL FIGURE 4

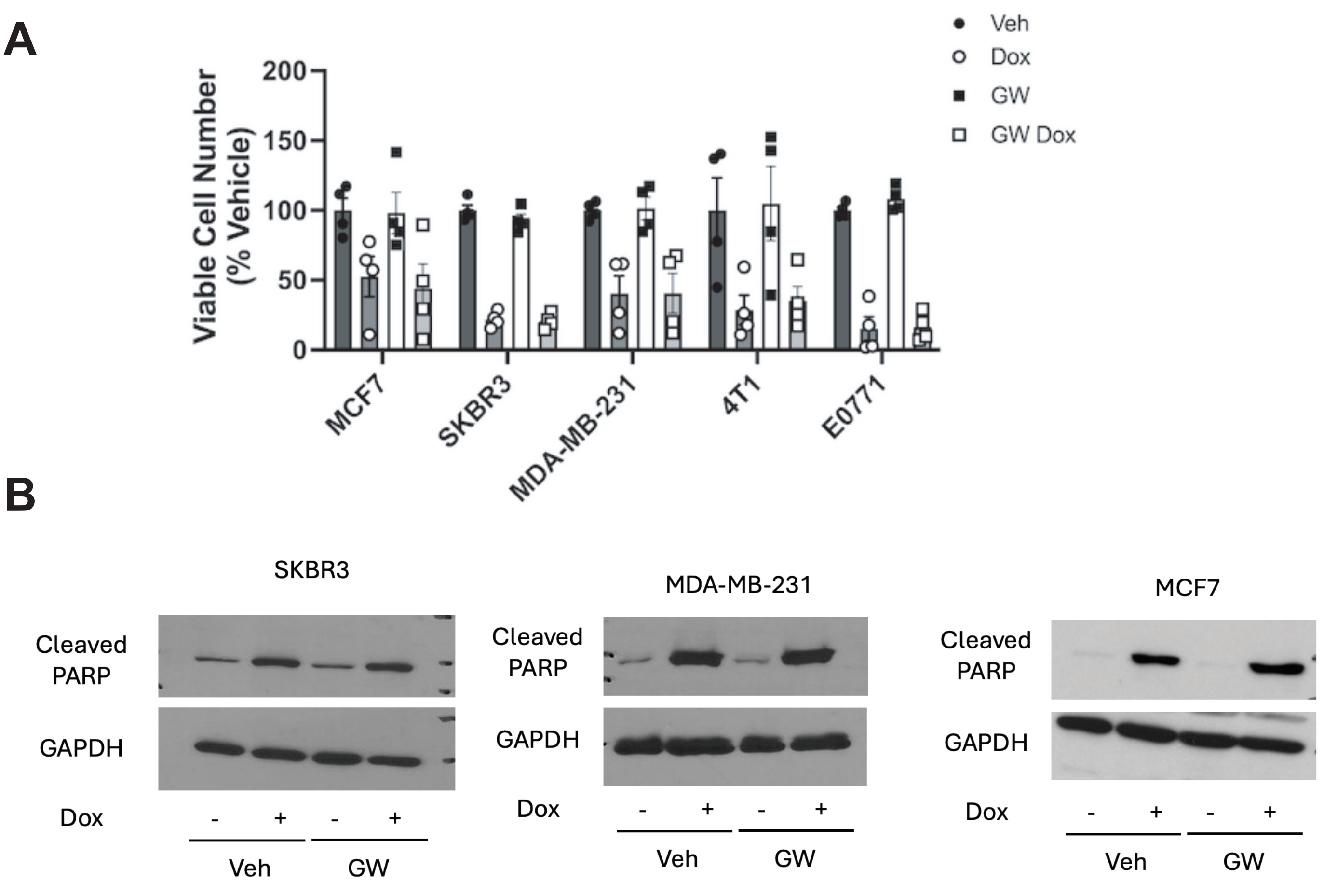


### SUPPLEMENTAL FIGURE 5

### SUPPLEMENTAL FIGURE 6

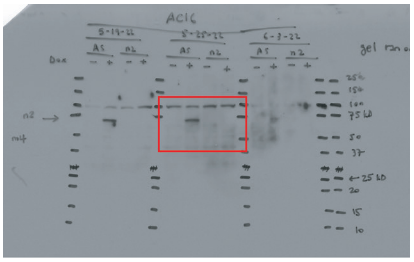

Figure 2E: nSMase2

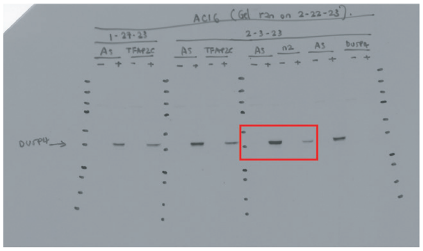

Figure 5C: DUSP4

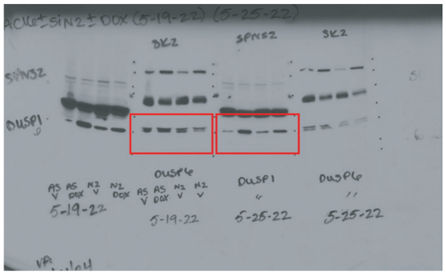

Figure 5C: DUSP6 and DUSP1

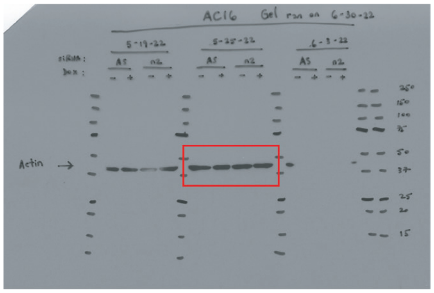

Figure 2E: Actin

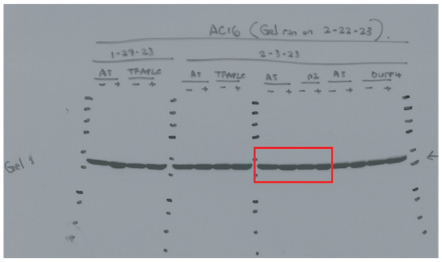

Figure 5C: Actin for DUSP4

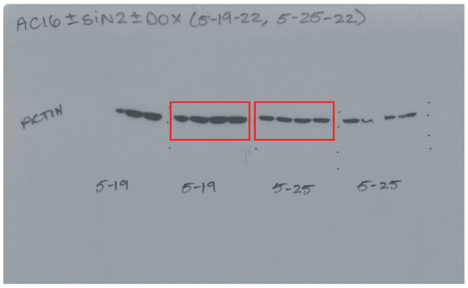

Figure 5C: Actin for DUSP6 and DUSP1

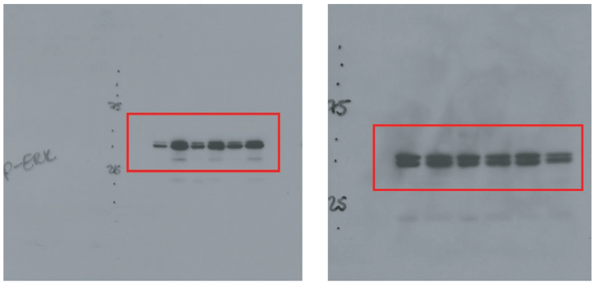


Figure 6E: p-ERK and ERK blots

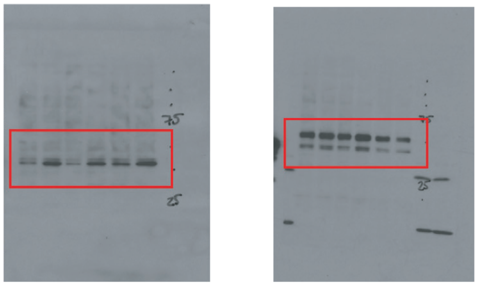


Figure 6E: p-JNK and JNK blots

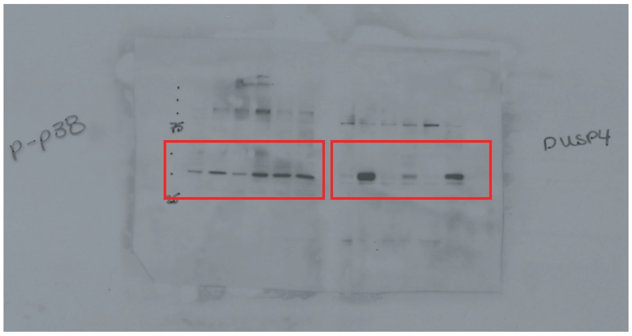


Figure 6E: p-p38 and DUSP4

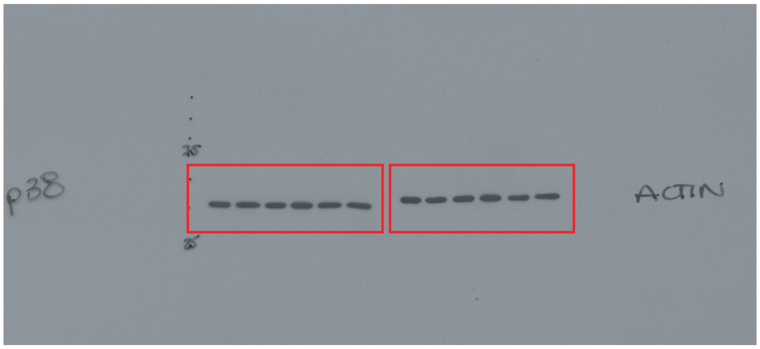


Figure 6E: p38 MAPK and Actin

| Gene Target | Company | Assay ID |
|---|---|---|
| AStar (negative) | Qiagen | 1027281 |
| SMPD3 (nSMase2) | Life Technologies | S30927 |
| p53 | Life Technologies | s605 |
| Top2A | Life Technologies | s14308 |
| Top2B | Life Technologies | s107 |
| DUSP4 | Life Technologies | s4373 |
| SMPD2 (nSMase1) | Life Technologies | s13170 |

**Supplemental Table 1: List of siRNA used**

| Target | Company | Catalogue # |
|---|---|---|
| nSMase2 | Santa Cruz | sc-67305 |
| DUSP1 | Cell Signaling | 48625S |
| DUSP4 | Cell Signaling | 5149S |
| DUSP6 | Cell Signaling | 50945S |
| Actin | Sigma | A2228 |
| phospho-ERK | Cell Signaling | 9101 |
| ERK | Cell Signaling | 4695 |
| phospho-JNK | Cell Signaling | 9251 |
| JNK | Cell Signaling | 9252 |
| phospho-p38 MAPK | Cell Signaling | 9211 |
| p38 MAPK | Cell Signaling | 9212 |
| P16/INK4a | Abcam | ab241543 |
| Cleaved PARP | Cell Signaling | 5174S |
| GAPDH | Cell Signaling | 5625S |

**Supplemental Table 2: List of antibodies used**

| Gene Target | Company | Assay ID |
|---|---|---|
| SMPD3 (human) | Life Technologies | Hs00920353_g1 |
| SMPD3 (mouse) | Life Technologies | Mm00491359_m1 |
| Top2A | Life Technologies | Hs01032137_m1 |
| Top2B | Life Technologies | Hs00172259_m1 |
| SMPD2 (nSMase1) | Life Technologies | Hs0090624_g1 |
| DUSP1 | Life Technologies | Hs00610256 |
| DUSP4 | Life Technologies | Hs01027785_m1 |
| Actin (human) | Life Technologies | Hs01060665_g1 |
| Actin (mouse) | Life Technologies | Mm02619580_g1 |

**Supplemental Table 3: List of Taqman primers used**

#### Supplemental Figure Legends

**Supplemental Figure 1. Doxorubicin induces comparable changes in sphingolipids in HL-1 cells.**

(A-C) HL-1 cardiomyocytes were treated with vehicle or Dox doses as shown for 24h. Lipids were extracted and analyzed by tandem LC/MS mass spectrometry for (A) Cer; (B) Sph and dhSph; (C) S1P and dhS1P. Data are expressed as mean ± SEM pmol/nmol lipid phosphate. (* $p<0.05$ ** $p<0.01$ *** $p<0.001$ $n = 4$ biological replicates).

(D) Lipid species in Dox-treated AC16 cells and Dox-treated hearts expressed as % vehicle controls calculated from data in Fig 1D (AC16) and Fig 1A (Heart) respectively.

**Supplemental Figure 2. nSMase2 is induced by Dox in HL-1 cells and iPSC-derived CMs.**

(A-E) The indicated cells were treated with vehicle or the Dox doses as shown for 24h. RNA was extracted, converted to cDNA, and gene expression analyzed by qRT-PCR for nSMase2 (A, B, E), or nSMase1 and nSMase2 (C, D) using actin as reference gene. Data are expressed as mean ± SEM of mean normalized expression (A, B) or mean ± SEM fold-change over vehicle (C,D) (*** $p < 0.01$, ** $p < 0.02$ vs vehicle; $n = 3$ biological replicates)

(F) Cells were treated with vehicle or the Dox dose shown for 24h. In vitro N-SMase activity was assayed as described in ‘Materials and Methods”. Data are expressed as mean ± SEM N-SMase activity. ($n = 4$ biological replicates)

(G-I) AC16 cells were pre-treated with negative control (si-AS, 20nM) or target siRNA (si-N2, 20nM) for 48h prior to treatment with vehicle (DMSO) or Dox (0.6μM) for 24h. Lipids were extracted and analyzed by tandem LC/MS mass spectrometry for (G) the Cer species shown; (H) Sph and dhSph; (I) S1P and dhS1P. Data are expressed as mean ± SEM pmol/nmol lipid phosphate. ($n = 4$ biological replicates).

**Supplemental Figure 3. Loss of nSMase2 is protective against Dox effects.**

(A) Plasma was collected from WT and fro/fro mice following chronic Dox treatment. Cardiac troponin levels were measured by ELISA. Data are expressed as mean ± SEM ng/ml.

(B) AC16 cells were pre-treated for 48h with negative control (si-AS, 20nM) or target siRNA (si-N2, 20nM) prior to treatment with vehicle (DMSO) or Dox (0.6μM, 24h). Viable cell number was assessed by MTT assay. Data are expressed mean ± SEM % fold-change over day 0. (n = 3 biological replicates)

(C) AC16 cells were pre-treated with negative control (si-AS, 20nM) or target siRNA (si-N2, 20nM) for 48h prior to treatment with vehicle (DMSO) or Dox (0.6μM) for 1h. Treatment was washed out and cells cultured in Vehicle or GW4869 for 72h before staining for senescence-associated beta-galactosidase activity, Pictures representative of at least three biological replicates.

(D) AC16 cells were pre-treated for 30 min with vehicle (DMSO:MSA) or GW4869 (10μM) prior to treatment with vehicle (DMSO) or Dox (0.6μM) for 1h. Treatment was washed out and cells cultured in Vehicle or GW4869 for 72h before staining for senescence-associated beta-galactosidase activity, Pictures representative of at least three biological replicates.

**Supplemental Figure 4. nSMase2 regulation of DUSPs is specific**

(A-C) Cells were pre-treated with negative control (si-AS, 20nM) or nSMase1 siRNA (siN1, 20nM) for 48h prior to treatment with vehicle (DMSO) or Dox (0.6μM, 24h). RNA was extracted, converted to cDNA, and gene expression for (A) nSMase1, (B) nSMase2, or (C) DUSP4 was assessed by qRT-PCR using actin as reference gene. Data are expressed as mean ± SEM of mean normalized expression (C) (*** $p < 0.01$, ** $p < 0.02$ vs vehicle. n = 4 biological replicates).

**Supplemental Figure 5. nSMase2 is dispensable for Dox-induced cell death in breast cancer cells.** (A, B) The breast cancer cell shown were pre-treated with GW4869 (10μM) or vehicle for 15 min prior to treatment with Dox (1μM) for 24h. (n = 4 biological replicates) (A) Viable cell number was assessed by MTT assay; (B) Protein was harvested and blotted for cleaved PARP and GAPDH as loading control.

**Supplemental Figure 6. Original immunoblots from manuscript.** Shown are the original western blot images taken for main figures. Red boxes highlight the selected area for the manuscript figures.
